# Engineering B cells to Express Fully Customizable Antibodies with Enhanced Fc Functions

**DOI:** 10.1101/2025.07.23.666432

**Authors:** Chun Huang, Atishay Mathur, Chan-Hua Chang, Xiaoli Huang, Hsu-Yu Chen, Zachary B. Davis, Karla O’Dell, Elizabeth A Shuman, Raymond W. Kung, Geoffrey L. Rogers, Paula M. Cannon

**Affiliations:** Department of Molecular Microbiology and Immunology, Keck School of Medicine of the University of Southern California, Los Angeles, California, USA; Department of Medicine, University of Minnesota, Minneapolis, Minnesota, USA; Department of Otolaryngology–Head & Neck Surgery, University of Southern California, Los Angeles, California, USA

## Abstract

Genome editing within the constant region of the immunoglobulin Heavy chain locus (*IGH*) can reprogram B cells to express Heavy chain only antibodies (HCAbs) containing custom antigen-recognition domains. HCAb-engineered cells express both surface B cell receptor (BCR) and secreted antibody isoforms and respond to antigen. By selecting alternate editing sites within *IGH*, we extended this approach to also allow customization of the constant (Fc) domain of the Heavy chain, producing HCAbs with enhanced effector functions or containing mutations to extend antibody half-life. We also introduced mutations to force obligate HCAb homodimers and prevent unwanted pairing with endogenous antibody chains. Finally, we showed that additional domains could be accommodated at the HCAb C-terminus and preferentially expressed in the secreted isoform. Together these data demonstrate the flexibility of the HCAb editing platform to express fully customized molecules that take advantage of the properties of B cells.

## Introduction

Monoclonal antibodies with either natural or engineered designs are effective treatments for many conditions^1–3^. However, when long-term administration is required, repeat infusions of these recombinant proteins can be costly and inconvenient. To address this limitation, we and others have proposed that B cells, including *ex vivo* differentiated plasma cells, could be engineered to serve as cellular factories for long-term secretion of therapeutic antibodies and other molecules *in vivo*^4–11^. In a further sophistication, inserting custom antibody components directly within the Immunoglobulin Heavy chain locus (*IGH*) can result in B cells that retain the ability to respond to a matched antigen^4–6,9,11^. In this way, *IGH*-engineered B cells have the potential to be a long-lived and boostable cell therapy.

We previously described a simplified approach to reprograming human B cells using CRISPR/Cas9 editing and homology-directed recombination (HDR) to insert custom antigen recognition domains (ARDs) into constant regions of the *IGH* locus^5^. By targeting insertion downstream of the CH1 exon in *IGHG1*, we produced antibodies that mimicked the design of the Heavy chain only antibodies (HCAbs) found in camelids, and which do not interact with antibody Light chains. Compared to strategies for engineering conventional antibodies^4,6,9,11^, which require replacement of variable domains from both Heavy and Light chains, the HCAb approach is much simpler and allows greater flexibility in the ARDs that can be accommodated. For example, we created HCAbs with ARDs based on both antibody (scFv and V_H_H) and non-antibody (CD4 protein) components^5^. We also confirmed that HCAbs could be expressed as both cell surface B cell receptors (BCR) and secreted antibody isoforms, depending on the differentiation state of the B cell, and that the engineered cells responded to antigen stimulation in a human tonsil organoid model of immunization^5^.

In the present study, we describe a further enhancement of HCAb engineering to allow the production of fully customizable HCAbs. By moving the site for ARD cassette insertion downstream within Heavy chain constant regions, we are additionally able to replace some or all of the exons comprising the Fc domain. We used this approach to introduce CH2 mutations that enhanced ADCC function and CH3 mutations to increase antibody half-life. Mutations in CH3 were also incorporated that resulted in obligate HCAb homodimers, thereby excluding unwanted interactions between HCAbs and endogenous Heavy chains in the same cell. Finally, by targeting insertion sites at the 3’ end of CH3, we were also able to incorporate additional protein domains at the C-terminus of the HCAbs, which could further enhance functionality. The alternate splicing mechanism that controls expression of BCR versus secreted antibody isoforms was preserved, and the C-terminal additions were preferentially expressed in secreted HCAbs, preventing potential interference with the function of the BCR. Together, these capabilities provide a B cell engineering platform with unprecedented flexibility to accommodate antibody and non-antibody designs that retain the characteristics of natural antibody production.

## Results

### Genome engineered HCAbs with custom CH2 domains

We previously described a genome editing approach to insert cassettes comprising a B cell specific promoter (EEK) and a custom ARD downstream of the CH1 exon in the constant region of *IGHG1.* This results in linkage of the ARD to the endogenous IgG1 Fc domain, and allows expression of both secreted and membrane-anchored isoforms of the resulting HCAb, depending on the differentiation state of the B cell^5^. Although this format provides great flexibility in the nature of the inserted ARD, an ability to also modify the Fc domain is desirable, since this region influences a range of antibody properties^12,13^. One approach to achieve this is to insert the ARD cassette into constant regions other than *IGHG1*, such as *IGHA*, which would result in HCAbs with properties determined by the corresponding Fc domain. However, to also take advantage of non-natural Fc designs, we expanded the editing approach to allow specific replacement of part or all of the Fc domain. An example of this concept is shown in **Figure 1a**, where the EEK-ARD cassette insertion is targeted to the intron downstream of the CH2 exon of *IGHG1*, and the cassette also includes the bypassed Hinge plus CH2 sequences.

**Figure 1.**
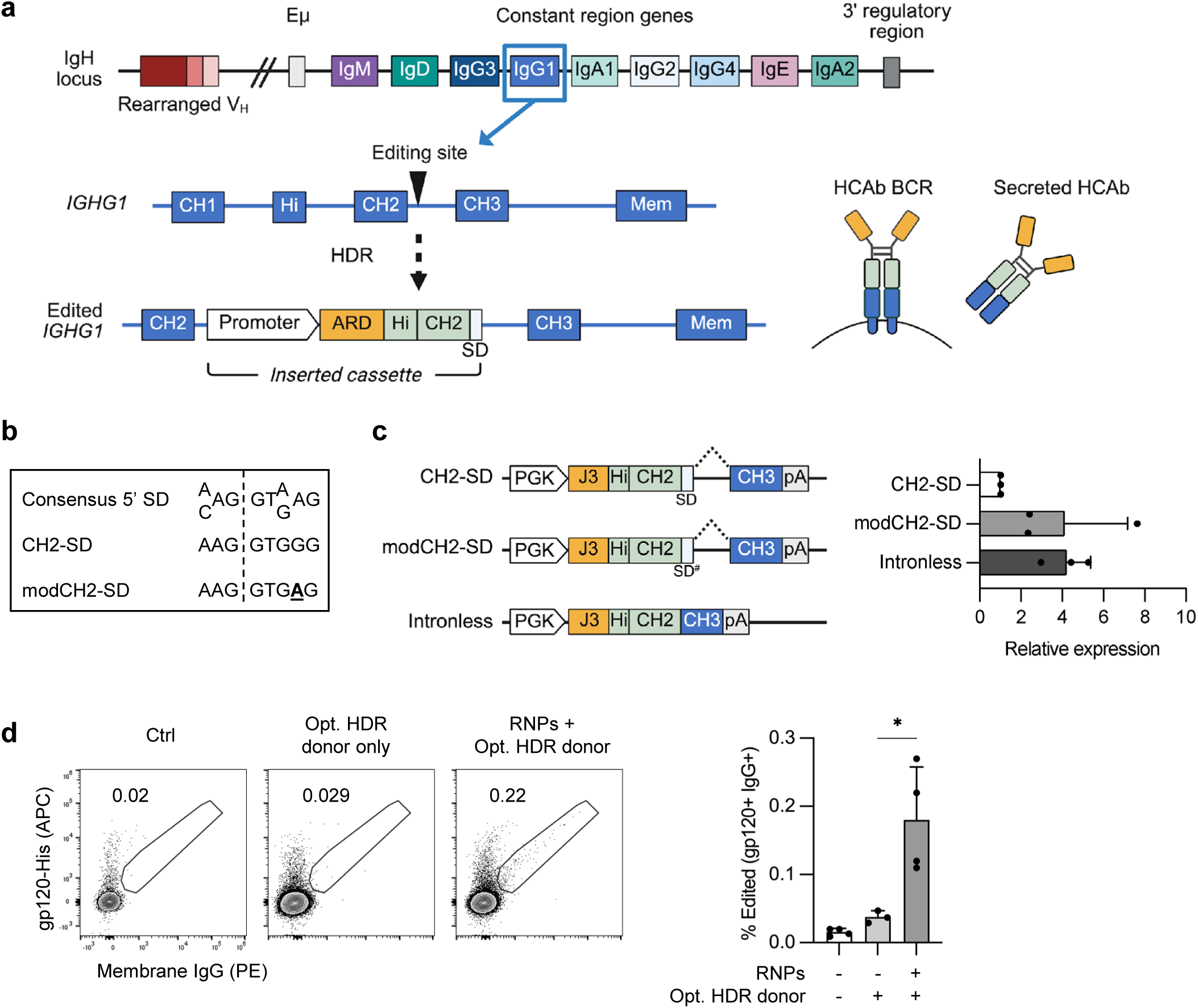
Genome editing to create CH2-modified HCAbs. (**a**) CRISPR/Cas9 and homology-directed repair (HDR) genome editing introduces a cassette comprising a B cell-specific promoter, antigen recognition domain (ARD) and codon wobbled Hinge and CH2 exons (light green) into the intron downstream of the *IGHG1* CH2 exon. Splicing to endogenous CH3 and membrane domains results in expression of either BCR or secreted HCAb isoforms. (**b**) The splice donor at the boundary of the CH2 exon and downstream intron (CH2-SD) was modified to more closely match the consensus 5’ SD. (**c**) Plasmids containing expression constructs mimicking the expected final gene-edited configurations with the original and modified CH2-SD were transfected into 293T cells. A plasmid without the CH2 intron or SD was used as a control to measure expression without splicing. Secretion of J3-CH2 HCAbs was measured by IgG ELISA and made relative to the unmodified CH2-SD construct (*n* = 3). (**d**) Raji cells were edited with CH2-g1 Cas9 RNPs plus the optimized plasmid HDR template (codon wobbled Hinge and CH2 exons, modCH2-SD). Editing was measured by flow cytometry at 7 days post-editing. A representative example and graph showing results from *n* = 4 independent experiments are shown. Error bars represent mean ± SEM. Statistical analysis was performed using unpaired 2-tailed t test. * *p* < 0.05.

We used this approach to create HCAbs with custom CH2 domains. Genome editing reagents were optimized for HDR insertion at the CH2 intron using methods previously described^5^. We identified a suitable guide RNA, CH2-g1, with good on-target activity at *IGHG1* but undetectable off-target activity at related *IGHG* loci, and which supported HDR-mediated insertion of short oligonucleotides at the target site in human Raji B cells (**Extended Fig. 1a-b; Table S1**). A complementary HDR donor plasmid was constructed containing the EEK promoter, an ARD based on the anti-HIV V_H_H domain J3^14,15^, and codon wobbled IgG1 Hinge and CH2 domains, flanked by sequences homologous to the endogenous *IGHG1* locus (**Table S2**). Codon wobbling of the IgG1 Hinge and CH2 domains in the cassette was necessary to minimize homology with the endogenous *IGHG1* sequences that otherwise interfered with the intended HDR edit (**Extended Fig. 1c-d**). However, despite molecular evidence of successful editing with the codon wobbled donor construct, the expected J3-CH2 BCR was not detected on the surface of the edited Raji B cells (**Extended Fig. 1e**).

We hypothesized that the lack of expression of the J3-CH2 BCR was due to impaired splicing. To test this, we modified the natural splice donor at the boundary between the CH2 exon and intron (CH2-SD) in the HDR donor to better match the consensus SD sequence^16,17^ (**Fig 1b**). Plasmid constructs mimicking the final expected gene-edited configurations were transfected into 293T cells and confirmed that J3-CH2 HCAb expression was enhanced by this SD modification (**Fig. 1c**). When used to edit Raji cells, an HDR template containing both the codon-wobbled IgG1 sequences and the modified CH2-SD now supported cell surface expression of the J3-CH2 BCR (**Fig. 1d**). Enriching the population of edited Raji cells by FACS allowed us to harvest J3-CH2 HCAbs from culture supernatants and confirm that they had the expected specific activity in an HIV neutralization assay (**Extended Fig 1f**).

### Functionality of CH2 HCAb edited primary B cells

Primary human B cells from PBMCs were activated and edited to express J3-CH2 HCAbs using Cas9 RNPs and AAV6-based homology donor vectors (**Fig. 2a**), as previously described^5^. Using this protocol, the editing efficiency reached about 40% (**Fig. 2b**). A switch from expression of BCRs to secreted antibody isoforms occurs as B cells differentiate into antibody-secreting cells (ASCs) and is a critical aspect of B cell biology. To confirm that the edited cells retained this activity, J3-CH2 HCAb edited cells were differentiated towards ASCs and analyzed by flow cytometry (**Extended Fig. 2**). We observed that the percentage of ASCs increased to a similar extent in gated HCAb+ sub-populations from edited samples compared to bulk CD19+ B cells from unedited samples, confirming that the edited cells retained a normal differentiation capability (**Fig. 2c**). We also observed a similar increase in the secretion of both total IgG and J3-CH2 HCAbs per cell (6-fold versus 8.4-fold) as the cells were differentiated into ASCs (**Fig. 2d**).

**Figure 2.**
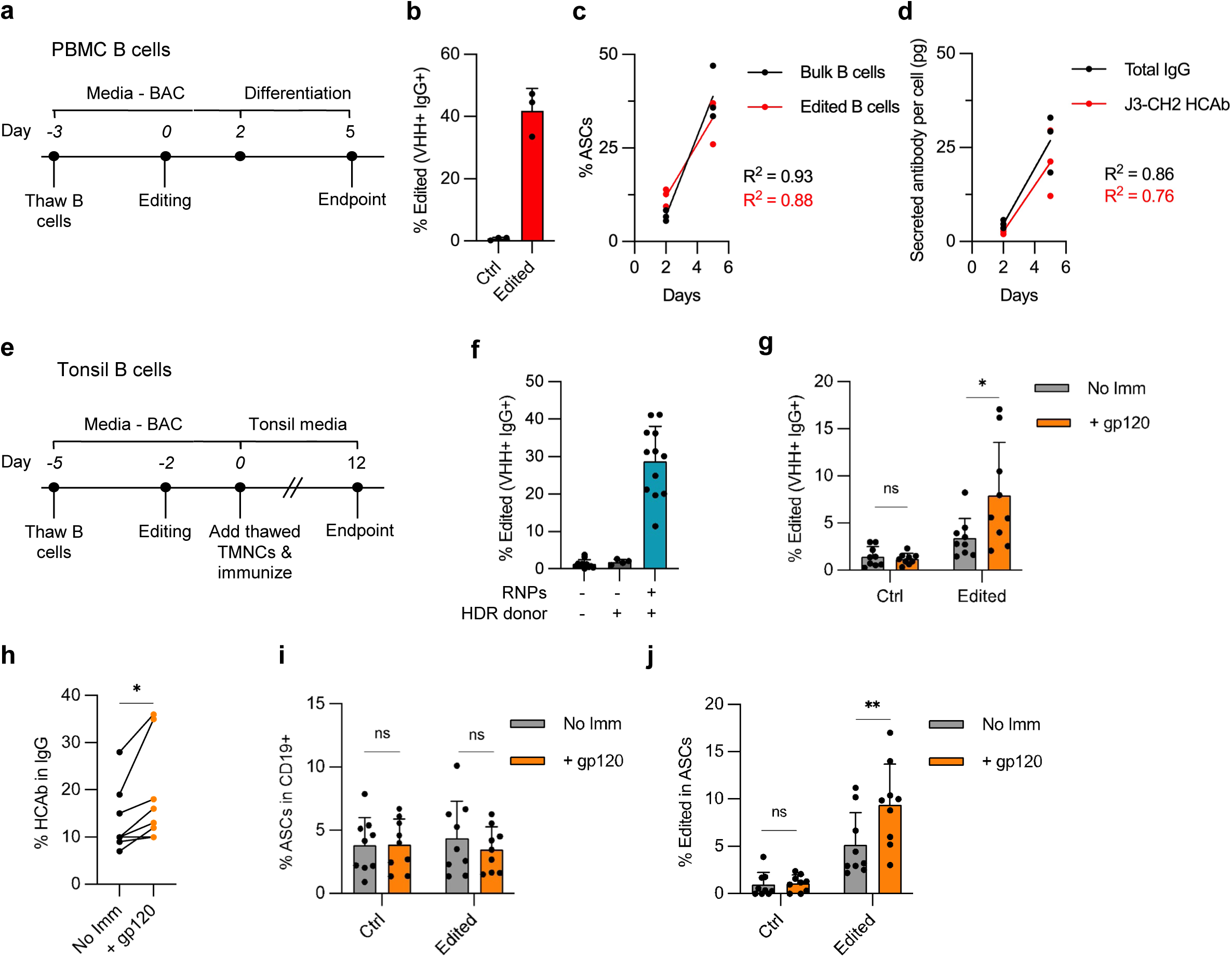
Functionality of CH2 HCAb edited primary human B cells. (**a**) Timeline for PBMC-derived B cell cultures. B cells were activated in BAC, edited using CH2-g1 RNPs and an AAV6 vector containing a J3-CH2 HCAb cassette (MOI = 1e4 vg/cell) on day 0, and cultured in differentiation media from days 2 to 5. (**b**) Percent edited (J3-CH2 HCAb+) B cells on day 2 by flow cytometry (n = 3, independent donors). (**c**) ASC differentiation days 2 to 5 in bulk CD19+ B cell populations from unedited samples and HCAb+ edited subset from edited samples, detected by flow cytometry. Gating strategy is shown in **Extended Fig. 2a**. Statistical analysis was simple linear regression with ANCOVA to compare regression lines, which were not significantly different (*p* = 0.88). (**d**) Supernatants were harvested from edited and control samples at days 2 and 5, and total IgG or J3-CH2 HCAbs measured by ELISA. Antibody secretion per cell was calculated for each timepoint. Statistical analysis was simple linear regression with ANCOVA to compare regression lines, which were not significantly different (*p* = 0.25). (**e**) Timeline for tonsil organoid cultures. Tonsil B cells were activated and edited as in (a), co-cultured with autologous tonsil mononuclear cells (TMNCs), and immunized with HIV gp120 plus Adju-phos adjuvant. Media were changed every 2-3 days and cells harvested at day 12 for analysis. (**f**) Percent edited (J3-CH2 HCAb+) tonsil B cells at day 0 (n=12 donors), prior to establishing tonsil co-cultures. (**g**) Response of control and edited tonsil organoid cultures (n=9 donors) at day 12 post-immunization (+gp120) or no immunization (No Imm.) A significant increase in J3-CH2 HCAb+ cells was seen in the edited population after immunization. See also **Extended Fig. 2b**. (**h**) Percentage of J3-CH2 HCAbs in total IgG in supernatant from matched co-cultures containing edited B cells at day 12, with and without immunization. See also **Extended Figs. 2c, d. (i**) No differences observed in percentage of ASCs in bulk CD19+ B cells from control or edited tonsil co-cultures, and with or without immunization, on day 12. (**j**) A significant increase was observed in percentage of J3-CH2 HCAb+ edited cells in the ASC population in the edited tonsil co-cultures after immunization. See also **Extended Fig. 2e**. Error bars represent mean ± SEM. The statistical analyses in (g-j) are paired 2-tailed t test between immunized and unimmunized samples. * *p* < 0.05, ** *p* < 0.01, ns not significant.

We next investigated whether J3-CH2 HCAb edited cells responded to a matched antigen. To do this we used a tonsil organoid model of immunization^18,19^ that we had previously adapted to assess HCAb-edited B cells^5^ (**Fig. 2e**). In brief, B cells were isolated from waste tonsil tissue and edited using the same protocol as for PBMC-derived B cells (**Fig. 2f**). Tonsil organoid cultures were established by combining a small number of J3-CH2 HCAb edited B cells with autologous tonsil mononuclear cells, resulting in a final frequency of 2-3% edited B cells. Following immunization with HIV gp120 antigen plus adjuvant, the frequency of the edited cells increased, and by day 12 was approximately 2.5-fold higher in co-cultures with, than without, immunization. (**Fig. 2g; Extended Fig. 2b**). This fold-expansion in response to antigen was similar to the rates observed in other studies for tonsil organoids treated with flu vaccine^18^ and agrees with our previous findings for J3 HCAb edited cells^5^. At the same time, the amount of secreted J3-CH2 HCAbs as a percentage of total IgG increased in the immunized edited co-culture, while the total secreted IgG did not change (**Fig. 2h, Extended Fig. 2c-d**). Finally, although the percentage of ASC in the total B cell population did not differ across any of the 4 experimental arms at day 12, the percentage of edited cells within the ASC population was significantly increased following immunization (**Fig. 2i,j; Extended Fig. 2e**). Together these data show that J3-CH2 HCAb edited cells responded to the gp120 antigen by clonal expansion, differentiation towards ASC, and secretion of J3-CH2 HCAbs.

### Modifying the CH2 domain to enhance ADCC function

The ability to alter the sequence of the CH2 domain opens up the possibility of introducing mutations to enhance effector functions, such as antibody-dependent cell cytotoxicity (ADCC). This occurs when antibodies bound to the surface of a target cell are recognized by Fc gamma receptors (FcγRs) on natural killer (NK) cells, resulting in NK activation and degranulation, and lysis of the targeted cells. Mutations that enhance ADCC have been identified in the CH2 domain, such as the GASDALIE (AE) combination that skews antibody binding towards activating FcγRs^20,21^. In contrast, the LALA-PG (LA) mutations nullify Fc function by reducing FcγR binding^22,23^.

We incorporated AE and LA mutations into J3-CH2 HCAb HDR templates and edited Raji cells as before (**Fig. 3a; Table S2**). Supernatants from sorted edited Raji cells were harvested and ADCC assays performed using HIV-infected cells as target cells. We included recombinant anti-HIV antibodies 3BNC117 and PGT121 as positive and negative controls, respectively^24–26^. Incubation of the HIV-infected target cells with NK-92 effector cells in the presence of the supernatants or antibodies allowed us to quantify target cell killing by flow cytometry (**Extended Fig. 3a**). As expected, the percentage of HIV-infected cells dropped significantly in samples incubated with the positive control antibody 3BNC117, but did not change in samples with the negative control antibody PGT121. For the non-mutated J3-CH2 HCAb supernatants, we also observed evidence of ADCC activity, which was significantly enhanced by the inclusion of the AE mutations, but was reduced to background levels by inclusion of the LA mutations.(**Fig. 3b; Extended Fig. 3b**). These data support that the CH2 HCAb genome editing strategy allows the introduction of desired CH2 domain mutations to modulate antibody effector functions.

**Figure 3.**
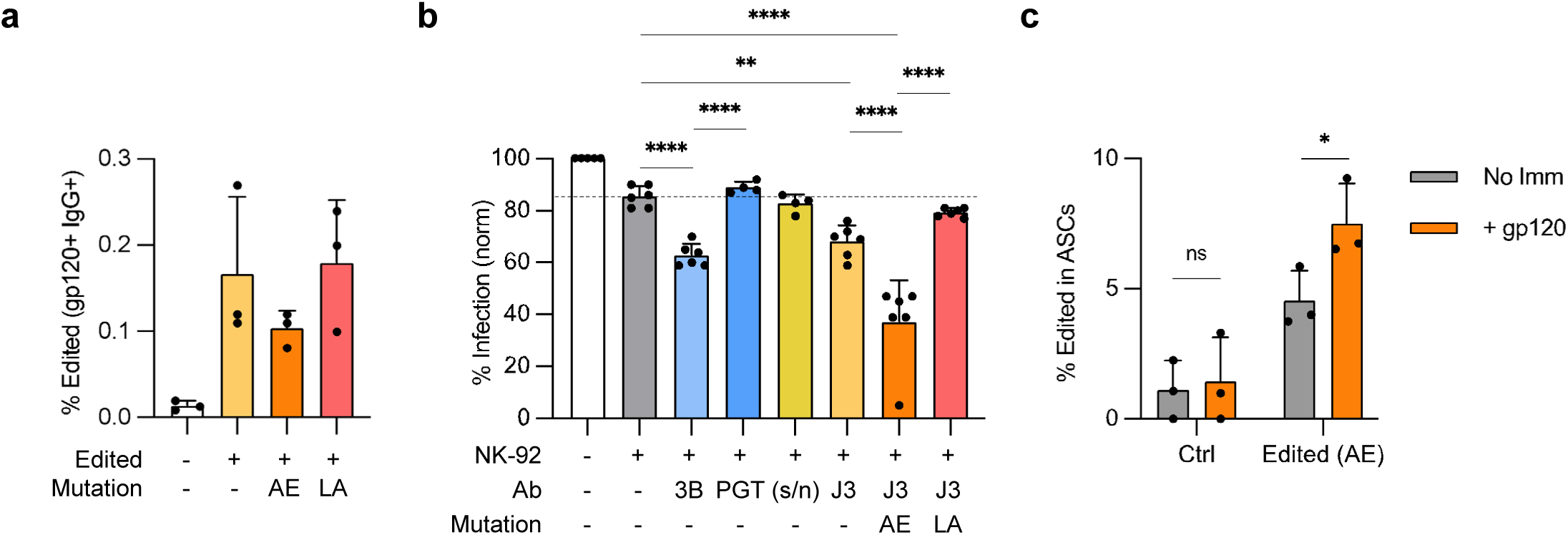
CH2 HCAb editing to incorporate ADCC mutations. (**a**) Raji cells were edited with CH2-g1 Cas9 RNPs and J3-CH2 plasmid HDR templates, including with enhancing (AE) or null (LA) ADCC mutations. Editing efficiency was measured day 7 post-editing as BCR+ cells by flow cytometry (*n* = 3). (**b**) ADCC activity was measured as a reduction in the percentage of HIV-infected cells due to NK-mediated cytotoxicity in presence of indicated antibodies (*n* = 3, with technical duplicates per trial). Recombinant antibodies 3BNC117 (3B) and PGT121 (PGT) were used as positive and negative controls, respectively. Supernatants were harvested from sorted J3-CH2 HCAb edited Raji cells (J3) containing the indicated mutations, or from control non-edited Raji cells (s/n). The reductions in HIV-infected cells were normalized between experiments to account for difference in initial HIV+ cell populations. Dotted line indicates background ADCC activity when incubated with NK-92 cells only (grey). (**c**) Control non-edited or J3-CH2(AE) HCAb edited tonsil B cells were incorporated into tonsil organoids, treated with or without gp120 immunization (*n* = 3 donors), and the percent J3-CH2 HCAb+ cells (VHH+, IgG+) in the ASC population measured at day 12. Error bars represent mean ± SEM. Statistical analysis in (b) was 1-way ANOVA with multiple comparisons, while (c) was a paired 2-tailed t test. * *p* < 0.05, ** *p* < 0.01, *** *p* < 0.001, **** *p* < 0.0001, ns not significant.

Finally, we used the tonsil organoid system to evaluate the response of J3-CH2(AE) HCAb edited B cells to immunization. This confirmed that the CH2 HCAb editing strategy and the inclusion of AE mutations in the CH2 domain did not alter the ability of the HCAb BCR to mediate an antigen-specific response, and that the cells could express both BCR and secreted isoforms (**Fig. 3c; Extended Fig. 3c-d**).

### Engineering fully customized HCAbs

We next adapted the editing strategy to target insertions to the end of the CH3 exon, which would allow for replacement of the whole Fc domain, and the introduction of mutations into CH3. Preserving the function of the CH3 splice donor was an important consideration here, since alternate splicing at this site regulates production of the BCR versus secreted antibody isoforms. For this purpose, we found that *IGHG1* was not an optimal editing target since the sequence near the CH3 SD is identical in *IGHG2*, which could promote off-target CRISPR/Cas9 activity (**Extended Fig. 4a**). Instead, we selected a site in *IGHG4* with a unique sequence and performed a screen to identify a suitable guide RNA (G4-g2) (**Fig. 4a; Extended Fig. 4b-d; Table S1**).

**Figure 4.**
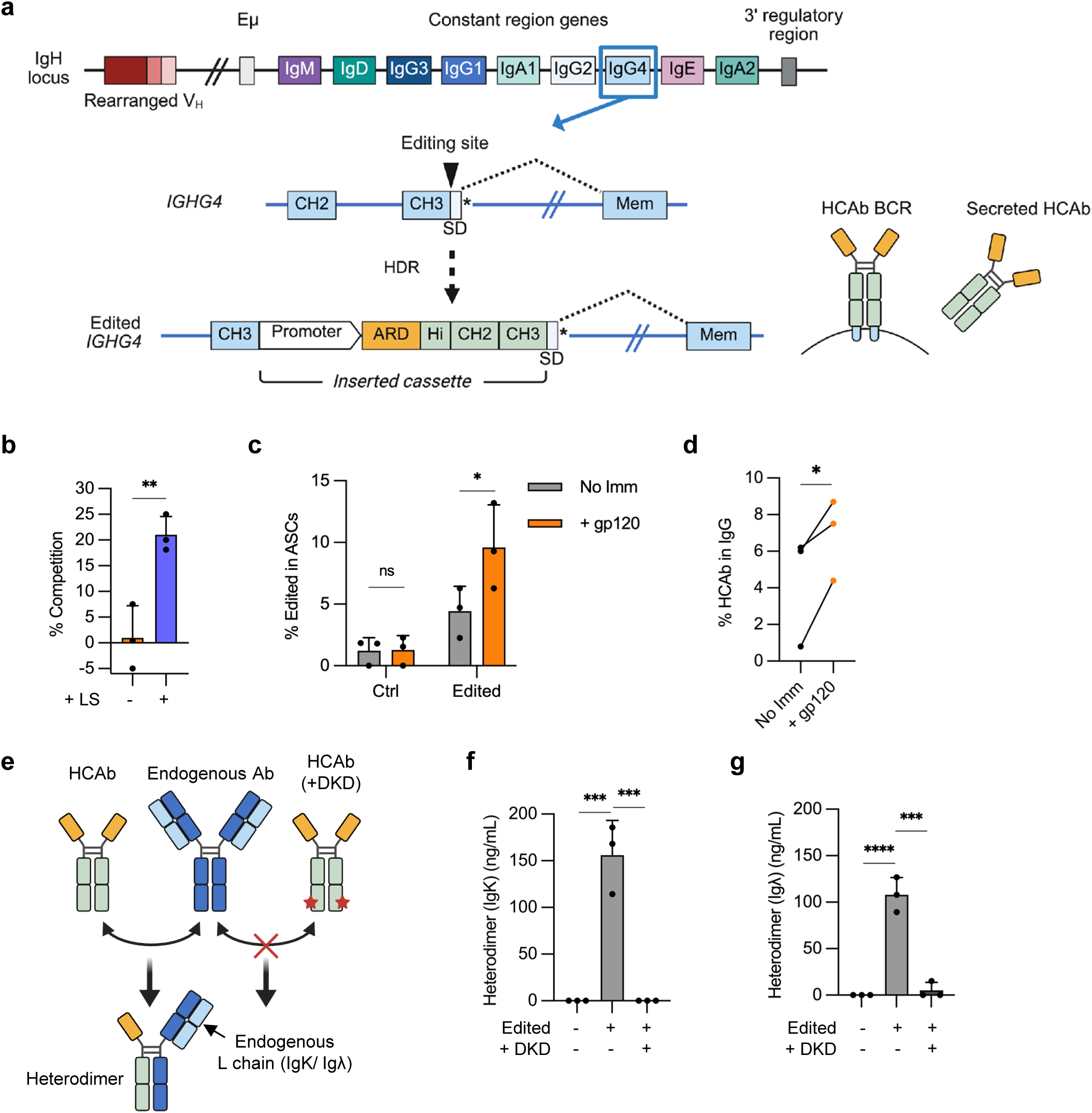
Genome editing for fully customized antibodies at *IGHG4*. (**a**) Editing approach, targeting the 3’ end of the *IGHG4* CH3 exon. The HDR template comprises a B cell-specific promoter, ARD, and codon-wobbled Hinge, CH2 and CH3 exons from *IGHG1*, which all provide customization possibilities. The cassette is inserted upstream from the SD in the CH3 exon and the stop codon (*) that produces the secreted antibody isoform. Following editing, the resulting HCAb contains an IgG1 Fc domain and can be expressed as either the BCR or secreted isoform through alternate splicing. (**b**) Supernatants from FACS-sorted edited Raji cells, with or without the LS mutations, were used in an FcRn competition binding assay (*n* = 3). (**c**) Tonsil B cells (*n* = 3 independent donors) were edited with G4-g2 Cas9 RNPs plus AAV6 HDR templates carrying a J3-IgG1 HCAb expression cassette, and both edited and control B cells were further cultured in tonsil organoids as described in Fig. 2e. Percent J3-IgG1 HCAb+ cells in the ASC population increased after gp120 immunization at day 12. (**d**) The percentage of J3-IgG1 HCAbs in total secreted IgG in supernatants of tonsil organoids containing edited cells increased at 12 days post-immunization compared to non-immunized cultures. (**e**) DKD mutations are predicted to prevent the formation of heterodimers between J3-IgG1 HCAb and endogenous IgG1, which are detected using gp120/Light chain ELISAs. (**f,g**) Tonsil B cells were edited to express J3-IgG1 HCAbs, with or without the DKD mutations. Supernatants (n=3) were harvested 2 days post-editing and heterodimers measured by gp120/IgK or gp120/Igλ ELISAs. Error bars represent mean ± SEM. Statistical analysis were performed by paired 2-tailed t test in (c), paired one-tailed t test in (d) and unpaired 2-tailed t test in (f, g). * *p* < 0.05, ** *p* < 0.01, *** *p*<0.001, **** *p*<0.0001.

As one potential application, the ability to engineer the CH3 domain could allow incorporation of mutations that extend antibody half-life, such as the LS mutations that promote interaction with the neonatal Fc receptor (FcRn) and enhance antibody recycling^27–29^. An HDR template was therefore designed to insert an expression cassette containing the EEK promoter, the J3 V_H_H ARD and a complete IgG1 Fc domain, with or without the LS mutations, at the *IGHG4* target site. As before, the template included codon-wobbled exons for the *IGHG1* Hinge, CH2 and CH3 domains, to reduce unwanted homology with endogenous sequences (**Table S2**). Raji cells edited at *IGHG4* to express J3-IgG1 HCAbs with the LS mutations produced HCAbs with the expected enhanced FcRn binding activity (**Fig. 4b; Extended Fig. 4e**).

We also confirmed that *IGHG4* CH3-exon edited primary B cells maintained the ability to respond to gp120 immunization in the tonsil organoid system and could express both BCR and secreted isoforms. Tonsil B cells were edited using G4-g2 Cas9 RNPs and AAV6 HDR vectors carrying a J3-IgG1 HCAb cassette, and tonsil organoids established with either edited or control B cells. Following immunization with gp120, the frequency of J3-IgG1 HCAb+ edited cells increased significantly in the ASC population (**Fig. 4c**), as did the percentage of J3-IgG1 HCAbs in the secreted IgG population (**Fig. 4d**).

Finally, we explored the possibility of including CH3 mutations to prevent the formation of heterodimers between engineered HCAbs and endogenous antibodies. Although HCAbs do not interact with Light chains, it is still possible for heterodimers to form between the Heavy chains in HCAbs and any endogenous Heavy chains expressed in the edited cells, creating unwanted bi-specific antibodies (**Figure 4e**). To prevent this, we introduced mutations into CH3 to promote homodimer pairing (K392D/D399K/K409D)^30^, here referred to as the DKD mutations (**Table S2**). We expressed both control and DKD versions of J3-IgG1 HCAbs from the *IGHG4* CH3 exon site in human tonsil B cells. Although we confirmed equivalent rates of editing by flow cytometry, we did note that the DKD variant of this HCAb resulted in lower levels of secretion than the unmodified version (**Extended Figure 4f-g**). Next, we assed heterodimer formation. Since HCAbs do not contain a Light chain partner, we can detect J3-containing heterodimers using gp120/Igκ or gp120/Igλ ELISAs^5^. This showed that inclusion of the DKD mutations eliminated the formation of heterodimers compared to the non-mutated J3-IgG1-HCAb (**Figure 4f-g**).

### Selective addition of C-terminal domain to secreted HCAbs

Appending domains to the C-terminus of antibodies is a common approach to expanding the functionality of recombinant antibodies. We therefore designed an editing strategy to incorporate C-terminal additions into the secreted form of an HCAb, while preserving the splicing switch from BCR to secreted isoform. We piloted this approach by adding GFP to the C-terminus of the J3-IgG1 HCAb cassette using the G4-g2 target site at the C-terminus of *IGHG4* CH3. The GFP domain was placed downstream of the natural CH3-SD, in a strategy designed to incorporate the domain into the secreted isoform but exclude it from the BCR (**Fig 5a**). Analysis of the GFP sequence identified a cryptic SD at the end of the coding sequence that was predicted to greatly inhibit the action of the natural CH3-SD^31^. We therefore mutated this GFP-SD to block its activity and at the same time modified the CH3-SD to be closer to the consensus SD sequence (**Fig. 5b**).

**Figure 5.**
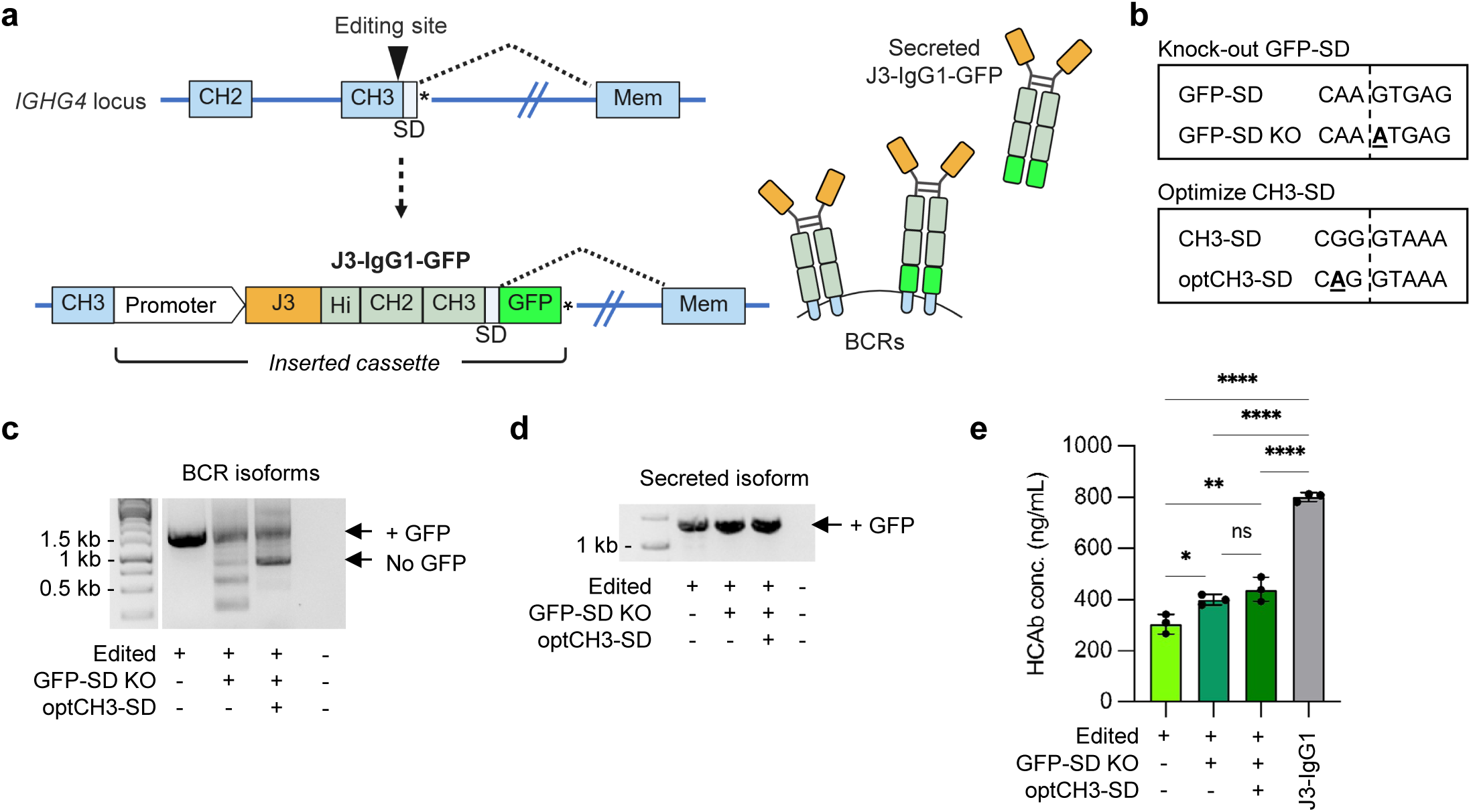
Addition of GFP domain to C-terminus of secreted HCAbs. (**a**) Editing strategy to create J3-IgG1 HCAbs with C-terminal GFP domain by editing at the G4-g2 target site in *IGHG4.* The GFP domain was placed downstream of the CH3-SD, to promote inclusion of GFP in the secreted HCAb isoform, followed by a stop codon (*). (**b**) Mutations to knock out a cryptic SD at the end of GFP and to optimize the CH3-SD. (**c,d**) mRNA from FACS-sorted J3-IgG1-GFP edited Raji cells, containing indicated SD mutations, was amplified by RT-PCR with primers specific for the BCR (c) or secreted HCAb (d) isoforms and analyzed by gel electrophoresis to identify transcripts of expected size, with and without a GFP domain. Identity of bands was further confirmed by Sanger sequencing (**Extended Fig. 5c, d**). (**e**) J3-IgG1-GFP HCAb secretion from FACS-sorted edited Raji cells, containing indicated SD mutations, was quantified by IgG ELISA. Supernatants from J3-IgG1 HCAb edited Raji cells (no GFP domain) served as a control (*n* = 3). Error bars represent mean ± SEM. Statistical analysis was performed by 1-way ANOVA with multiple comparisons: * *p* < 0.05, ** *p* < 0.01, *** *p*<0.001, **** *p*<0.0001, ns not significant.

Analysis of mRNA transcripts from edited Raji cells revealed that, as predicted, the GFP-SD in an unmodified GFP construct interfered with BCR splicing, resulting in bypassing of the CH3-SD and incorporation of the GFP domain into the BCR proteins (**Fig. 5c-d; Extended Fig.5a-d**). However, inclusion of the two SD modifications increased expression of the non-GFP BCR species (**Fig. 5c**) and also increased secretion of the J3-IgG1-GFP HCAb (**Fig. 5e**). Together, these data demonstrate that addition of a large domain at the C-terminus of a secreted HCAb is possible, but that elimination of cryptic SD sequences may be required to optimize BCR and HCAb production.

### Expression of the broadly-acting HIV inhibitor, eCD4-Ig

The HIV-1 entry inhibitor, eCD4-Ig, is an antibody-like molecule that interferes with binding of the virus to both the CD4 receptor and the CCR5 co-receptor molecules^32–34^. This results in broad neutralizing activity, making the molecule a candidate as a long-lasting HIV inhibitor. To express an HCAb that recapitulates the structure of eCD4-Ig, we used an ARD comprising the two N-terminal domains of CD4, and placed a peptide mimetic of CCR5 (R5m) as a C-terminal extension, with an IgG1 Fc to link these two anti-HIV domains (**Fig 6a**). Raji cells were edited to express the eCD4-IgG1 HCAb by targeting the C-terminus of *IGHG4* CH3, and editing was confirmed by flow cytometry (**Extended Fig. 5e**). RT-PCR and sequencing analyses indicated that both BCR and secreted isoforms were produced, with the R5m peptide present exclusively in the secreted isoform (**Fig. 6b; Extended Fig. 5f**).

**Figure 6.**
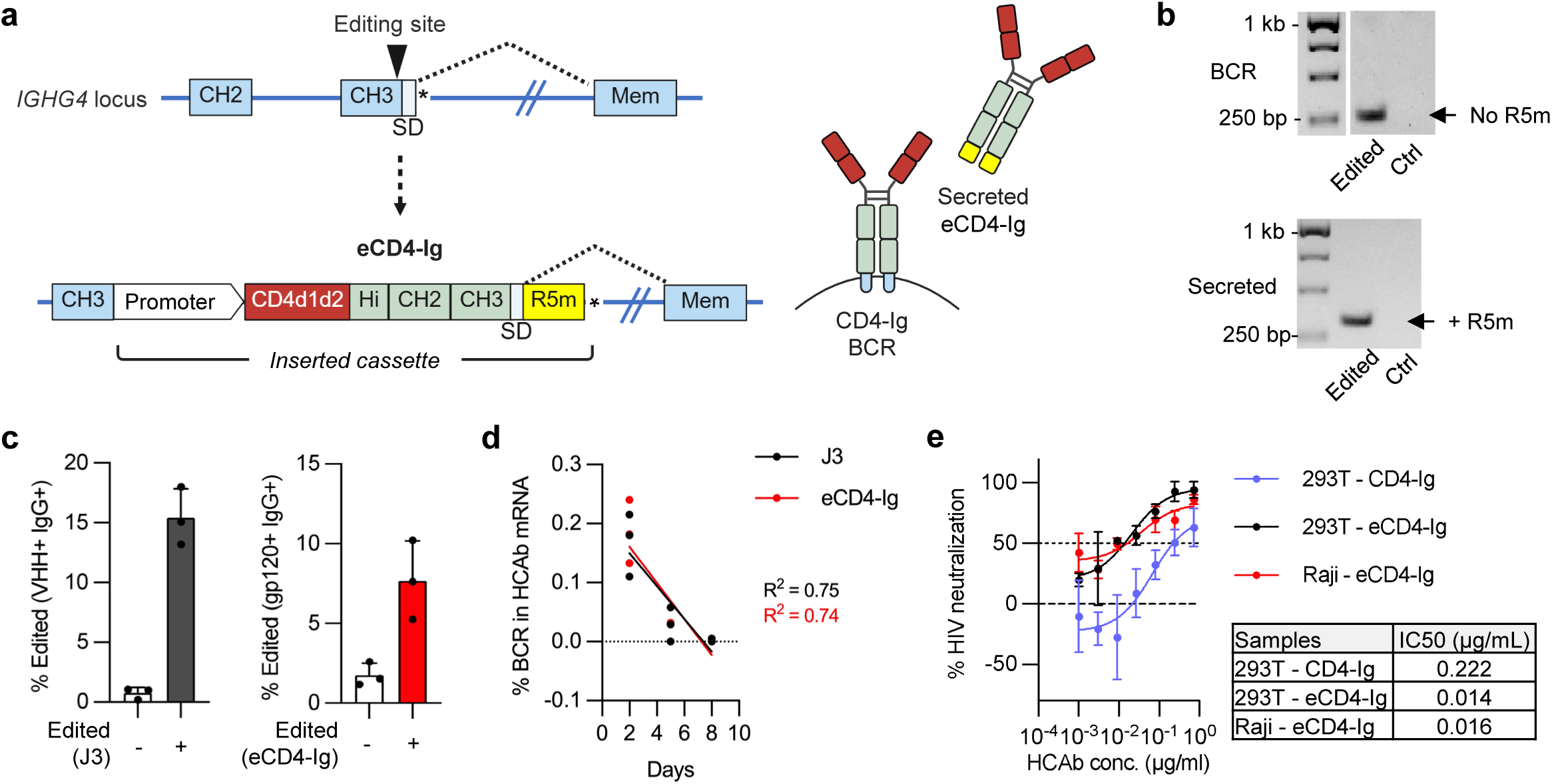
Secretion of eCD4-Ig from edited B cells. (**a**) Editing strategy to secrete eCD4-Ig from B cells. The inserted cassette comprises two domains of CD4 as the ARD and a CCR5 mimetic peptide (R5m) at the C-terminus, located downstream of CH3-SD to promote inclusion in the secreted HCAb isoform. (**b**) Raji cells were electroporated with G4-g2 Cas9 RNPs plus a plasmid HDR template for the eCD4-Ig cassette. mRNA from control or edited cells was used to amplify BCR or secreted HCAb species using specific primers. Only the secreted HCAbs had a size indicating inclusion of the R5m peptide. Identity of the bands was confirmed by Sanger sequencing (**Extended Fig. 5f**). (**c**) Human PBMC B cells were edited with G4-g2 Cas9 RNPs plus AAV6 HDR templates carrying either a control J3-IgG1 HCAb cassette, or an eCD4-Ig cassette and editing efficiency calculated by flow cytometry 2 days post-editing (*n* = 3). Gating strategy for eCD4-Ig is shown in **Extended Fig. 5g**. (**d**) Control and edited PBMCs were differentiated to ASCs, and the percentage of BCR transcripts in total HCAb mRNA species was quantified at indicated timepoints by reverse transcription and ddPCR. Simple linear regression was used to analyze the percentage BCR changes over time, and comparison of the regression lines for J3-IgG1 HCAb and eCD4-Ig samples showed no significant difference (p = 0.894). (**e**) HIV-1 neutralizing activity of eCD4-Ig secreted from FACS-sorted edited Raji cells was compared to recombinant eCD4-Ig and CD4-Ig (no R5m peptide) produced from transfected 293T cells (*n* = 2) and IC50 values were calculated from the curves.

Next, we addressed whether the C-terminal R5m addition had any impact on isoform switching. Both eCD4-IgG1 and control J3-IgG1 HCAb constructs were inserted into the *IGHG4* CH3 exon in human PBMC B cells (**Fig. 6c; Extended Fig. 5g**) and the cells differentiated towards ASCs, as previously described (**Fig 2a**). mRNA was harvested on days 2, 5 and 8 in culture, and used to quantify the percentage of BCR transcripts in the total HCAb mRNA. As expected, BCR isoforms decreased during ASC differentiation, becoming undetectable by day 8 (**Fig 6d**), and no differences were observed between the J3-IgG1 and eCD4-IgG1 transcripts. This indicates that inclusion of a C-terminal R5m domain downstream from CH3-SD did not impact isoform switching during B cell differentiation.

Finally, supernatants from FACS-sorted edited Raji cells were used in an HIV neutralization assay, which can distinguish the activity of proteins with (eCD4-IgG) and without (CD4-Ig) the C-terminal R5m peptide^33^. This revealed that the eCD4-IgG1 HCAbs secreted from Raji cells had a specific neutralizing activity most similar to the recombinant eCD4-Ig molecule containing R5m (**Fig. 6e**), confirming that the edited Raji cells secreted authentic eCD4-IgG1.

## Discussion

B cells are an emerging class of immune cell therapies^35^, due both to their potential to serve as *in vivo* factories for protein replacement, and because BCR engineering can be used to program their antigen specificity, in a manner analogous to engineered CAR/TCR T cells. Because of the complexity of antibody design, a common BCR engineering approach has been to insert a cassette comprising both a complete Light chain and a variable Heavy chain (VH) domain into the *IgH* locus, at locations between the VDJ region and the alternate constant region genes^4,6,11,36^. A consequence of this approach is that the Fc domain of the resulting BCR and antibody is provided by endogenous sequences, with the specific isotype determined by the natural process of class switch recombination. While this means that engineered cells can potentially diversify effector functions across multiple antibody isotypes, it does not allow control over the specific antibody isotype that is expressed or could evolve *in vivo.* Moreover, the antibodies produced are limited to naturally occurring constant regions and cannot take advantage of any of the enhanced designs available with recombinant antibody therapeutics^22^. In contrast, HCAb editing offers the flexibility to express antibodies with a significantly broader range of antigen recognition domains (ARDs) beyond traditional VH plus VL combinations^5^ and, as the present study shows, to link them to defined and fully customized Fc regions, all while maintaining functionality and antigen-specific B cell responses.

Using this expanded HCAb editing platform, we demonstrated the feasibility of introducing specific synthetic Fc region enhancements. Human B cells were reprogrammed to express antibodies with enhanced ADCC activity or lacking ADCC activity, or with mutations to extend their half-life. In an approach to improve the safety of HCAb engineering, we also engineered obligate homodimer constant regions that prevented cross-pairing between HCAbs and endogenous IgG1 Heavy chains. Combined with the lack of Light chain partners in the HCAb design, this ensures that no mixed antibody species are created in HCAb-edited cells, even if endogenous antibody chains are also co-expressed. The elimination of unwanted antibody chain pairing is a further advantage of HCAb engineering that is not easily available for approaches that maintain a Light chain partner and rely on endogenous Fc domains^35^.

We also demonstrated that non-antibody sequences could be added to the C-terminus of HCAbs and preferentially included in secreted isoforms. This allowed us to express the HIV entry inhibitor eCD4-Ig from engineered B cells, in an approach that exploited two of the unique features of HCAbs – the use of a non VH+VL ARD based on the N-terminal domains of CD4, and the placement of a CCR5 mimetic peptide at the C-terminus of the molecule, where it has been shown to function most optimally^32^. By allowing expression of eCD4-Ig as a BCR, these engineered B cells are expected to be responsive to the presence of the HIV gp120 antigen. This, in turn, suggests utility as a cell therapy to control HIV that could be responsive to the rebound viremia that occurs if antiretroviral therapy is suboptimal or withdrawn^37–39^.

Antibodies with larger C-terminal additions are also being developed as therapeutics, for example as immunocytokines^40^ or to recruit T cells or NK cells to cancer cells^41^. Towards that goal we demonstrated that the much larger GFP domain could also be expressed as a C terminal addition to a secreted antibody. Interestingly, while the CCR5 mimetic peptide was completely omitted from the BCR isoform, we observed incomplete exclusion of the GFP domain, in large part due to a cryptic SD at the end of the GFP sequence. This suggests that sequences appended at the HCAb C-terminus should be carefully screened for cryptic SD sequences, and that additional optimizations may be necessary^42–44^.

It is anticipated that a major application of engineered B cells will be to express antibodies that cannot easily be elicited through vaccination. These could include molecules with ARDs that provide broad protection against highly mutagenic viruses such as HIV and influenza, or that recognize the self-antigens that are targeted in certain cancer and autoimmune disease therapies. HCAbs further expand these possibilities by supporting non-antibody ARD components, since the exclusion of the Light chain partner means there is no requirement to maintain a natural V_H_ + V_L_ design. In addition, an increasingly important component of antibody drug design is optimization of the Fc domain. For example, anti-PD-1 checkpoint inhibitors or anti-CD40 antibodies designed to stimulate the immune system are produced with low effector function subclasses (typically IgG2 or IgG4) to prevent killing of T cells, or fatal inflammatory syndromes, respectively^45–47^. Conversely, some tumor killing antibodies have been designed with biologically enhanced Fc effector functions^48^. The flexibility of Fc design in HCAbs, together with the possibility of adding C-terminal domains, allows for fully synthetic designs to be imagined.

Although insertion of an ARD within the *IgH* constant region is a non-natural design, we demonstrated that HCAbs expressed from both IgG1 and IgG4 maintained appropriate regulation of BCR/antibody expression. We similarly expect that insertions at other constant regions will also be functional. In addition to this providing a method to select specific antibody isotypes, the ability to target alternate constant region genes could also support allow expression of more than one HCAb from a single cell. This could allow for combinations of HCAbs with different antigen specificities, or provide complementary effector subclasses, such as IgG for circulating activity and IgA for activity at mucosal surfaces.

The design features of HCAb engineering do raise a potential concern about the longevity of engineered cells. Since constant region sequences are excised from the genome during class-switch recombination, it is possible that an inserted HCAb cassette could be lost in this way. However, we have previously shown that this is not inevitable, as even some IgA^+^ B cells could be engineered at the upstream IgG1 locus, likely by insertion into an inactive *IGH* allele that was not subject to such recombination^5^. In this respect, engineering at IgG4 should be more advantageous than IgG1, as only the relatively rare IgE and IgA2 constant regions are downstream of IgG4. Finally, because our cells are engineered to respond to antigen, we expect that the selective pressure of the germinal center reaction will maintain the population of HCAb engineered cells, even if some cells do end up losing the cassette.

In sum, we have expanded the HCAb editing platform to allow for complete customization of antibody sequences, including the Fc domain, while maintaining the ability to respond to antigen. To our knowledge, the HCAb editing technology is the only B cell engineering approach described so far that allows for both the invariant expression of an advantageous Fc region and the synthetic modification of the Fc component, while maintaining the B cell’s ability to respond to the cognate antigen. We anticipate that this technology will be highly flexible and facilitate engineered B cell therapies for a broad scope of applications.

## Material and methods

### Genome editing reagents

Truecut v2 Cas9 protein was obtained from Thermo Fisher Scientific (Waltham, MA). Guide RNAs (gRNAs) targeting human *IGHG1* and *IGHG4* (**Table S1**) were designed by ChopChop (http://chopchop.cbu.uib.no) and CRISPOR (http://crispor.tefor.net) and synthesized with 2’-O-methyl 3’ phosphorothioate modifications (Synthego, Silicon Valley, CA). Single-strand oligonucleotide (ssODN) HDR templates to test each gRNA were designed as previously described^49^ and obtained from IDT (Coralville, IA) (**Table S1**). Plasmid HDR templates to insert HCAb expression cassettes were designed with symmetrical 750-bp homology arms flanking the gRNA target sites, synthesized as gBlocks (IDT) and cloned into the pAAV plasmid backbone (Cell Biolabs, San Diego, CA) with Infusion cloning (Takara Bio, Shiga, Japan) (**Table S2**). HCAb components included the J3 anti-HIV VHH domain^5^ and both the CD4 domains 1 and 2 and a CCR5 mimetic peptide from plasmid CMVR-eCD4-IgG2-v26^33,34^, a gift from Michael Farzan (Harvard University). The plasmid HDR templates were also used to synthesize AAV6 vectors by Vector Biolabs (Malvern, PA) or Charles River Laboratories (Wilmington, MA), and titered as previously described^50^.

### Cell line culture and genome editing

Raji cells (American Type Culture Collection (ATCC), Manassas, VA) and CEM.NKR.CCR5 cells (NIH HIV Reagent Program, BEI Resources) were cultured in complete RPMI (cRPMI): RPMI-1640 medium (VWR, Radnor, PA) supplemented with 10% fetal bovine serum (FBS) and 1% penicillin/streptomycin (P/S) (VWR). CD16+ NK92 cells^51^ were provided by Drs. Jeffrey Miller and Zachary Davis (University of Minnesota) and were cultured in MEMα medium (Gibco, Grand Island, NY) supplemented with 0.1mM 2-mercaptoethanol (Gibco), 500U/mL recombinant IL-2 (PeproTech, Cranbury, NJ), 12.5% heat-inactivated horse serum (Gibco), 12.5% heat-inactivated FBS, and 1% P/S. 293T cells (ATCC) were cultured in complete DMEM (cDMEM): DMEM (VWR) with 10% FBS and 1% P/S. All cell lines were maintained at 37°C and 5% CO_2_ and passaged 2 to 3 times a week.

Raji cells were genome edited by electroporation of gRNA/Cas9 RNPs using a 4D-X Nucleofector (Lonza, Basel, Switzerland) and the SG Cell Line 96-well Nucleofector® Kit (Lonza), per manufacturer’s instructions. In brief, Raji cells were washed with phosphate buffered saline (PBS) and resuspended in SG buffer at 2×10^7^ cells/mL, mixed with pre-complexed RNPs (3 µg Cas9 protein and 60 pmol gRNA), and 1.4 μg plasmid or 100 pmol ssODN HDR templates. Mixtures were loaded into 16-well Nucleocuvette™ Strips (Lonza) and electroporated using pulse code DS-104.

### Primary B cell culture and genome editing

Frozen peripheral blood CD19+ B cells (STEMCELL Technologies, Vancouver, BC, C) were cultured and edited as previously described^5^. In brief, B cells were thawed and cultured at 5×10^5^ cells/mL in XF/BAC media, Immunocult-XF T cell expansion media (STEMCELL Technologies) supplemented with 1% penicillin/streptomycin/amphotericin B (PSA) (Lonza) and 2% Immunocult-ACF Human B Cell Expansion Supplement (BAC, STEMCELL Technologies. After 3 days of culture, B cells were resuspended in XF media at a concentration of 5×10^7^ cells/mL and 10 μL of cells incubated with AAV6 vectors (MOI = 1×10^4^) in a U-bottom 96-well plate at 37°C and 5% CO_2_ for 1 hour. This was then mixed with 90 μL BTXpress Electroporation Buffer (BTX, Holliston, MA) plus pre-complexed Cas9 RNPs (15 µg Cas9 protein and 300 pmol gRNA) and electroporated in 2 mm gap electroporation cuvettes (BTX) with a BTX ECM 830 machine at 250 V for 5 ms. B cells were then cultured in XF/BAC media at a density of 5×10^5^ cells/mL for 2 days.

B cells were differentiated as previously described^5^. In brief, after culturing in XF/BAC for 2 days post-electroporation, B cells were cultured in plasmablast generation media (XF media supplemented with 10 ng/mL IL-2 (R&D Systems, Minneapolis, MN), and 50 ng/mL IL-6, 50 ng/mL IL-10 and 10 ng/mL IL-15 (all from PeproTech), and 1% PSA), for 3 days. Then, cells were cultured in plasma cell generation media (XF media supplemented with 50 ng/mL IL-6, 10 ng/mL IL-15, plus 500 U/mL IFN-α (R&D Systems), and 1% PSA) for an additional 3 days.

### Tonsil organoids

Tonsil tissue was received from the Department of Otolaryngology-Head & Neck Surgery, Keck School of Medicine of the University of Southern California (USC), as anonymous waste samples, approved by USC’s Institutional Review Board (protocol HS-17-00023-AM001). Tonsil mononuclear cells (TMNCs) were harvested as previously described^5^. In brief, tonsil tissues were dissected into small pieces, mashed through 70 µm cell strainers, and isolated by Ficoll density gradient separation (700 x g, 20 min, no brake). A portion of TMNCs was used to purify B cells with the EasySep Human CD19 Positive Selection Kit II (STEMCELL Technologies) per the manufacturer’s instructions. TMNCs and purified CD19+ B cells were frozen in Cryostor CS-10 (STEMCELL technologies) and stored in liquid nitrogen until use.

Tonsil B cells were thawed, cultured, and edited as described above for peripheral blood CD19+ B cells. Two days post-electroporation, TMNCs from the matched donor were thawed and mixed with edited or unedited (control) B cells at a 19:1 ratio. Cells were resuspended in tonsil culture media, comprising RPMI with GlutaMAX (Gibco) supplemented with 10% heat-inactivated FBS, 1% P/S, 1% MEM Non-Essential Amino Acids Solution (Gibco), 1 mM Sodium Pyruvate (Thermo Fisher), 100 µg/mL Normocin (InvivoGen, San Diego, CA), 1% Insulin-Transferrin-Selenium (Gibco), and 200 ng/mL recombinant human B cell-activating factor (BioLegend, San Diego, CA). The cells were resuspended to a final concentration of 6×10^7^ cells/mL for 24-well transwell plates (Millipore-Sigma, Burlington, MA) or 2×10^7^ cells/mL for 96-well transwell plates (Millipore-Sigma) and 100 μL of the cell mixture was added into the upper well of the transwell, with tonsil culture media alone added to the basolateral chamber. Media in the basolateral chamber was changed every 2 to 3 days until cells were harvested.

For immunization, 1 μg (24-well) or 0.33 μg (96-well) of recombinant HIV-1 JR-CSF gp120 (Immune Technology, New York, NY) was mixed with Adju-Phos (InvivoGen) following the manufacturer’s instructions. Antigen/adjuvant mixture was added to upper chamber immediately after plating cells.

### Genome editing analysis

Edited or unedited (control) cells were pelleted and lysed in QuickExtract™ DNA Extraction Solution (Biosearch Tech, Hoddesdon, UK) following the manufacture’s protocol. One μL of cell lysate was used for each PCR reaction with AmpliTaq Gold 360 Master Mix (Thermo Fisher) plus primer sets to amplify the region around predicted gRNA cut sites, both on and off-target (**Table S3**). PCR reactions were run on a C1000 Touch Thermal Cycler (Bio-Rad, San Francisco, CA). PCR products were subject to Sanger Sequencing (Azenta, Burlington, MA) and analyzed by ICE assay [https://ice.synthego.com/<u>#/</u>] (Synthego Performance Analysis, ICE Analysis). Data was used to calculate the percentage of different outcomes (e.g. % indels, % knock out or % small fragment edited outcomes).

### Nested in-out PCR

Cells were pelleted, and genomic DNA was purified using the DNeasy Blood & Tissue Kit (Qiagen, Hilden, Germany). Fifty nanograms of genomic DNA was used for each PCR reaction, employing AmpliTaq Gold 360 Master Mix along with a primer set designed to amplify the CH2 intron edited regions. This primer set comprised a forward (Fwd) primer binding upstream from the L-HA and a reverse (Rev) primer targeting the EEK sequence (**Table S3**). Subsequently, the PCR product was diluted 1:50 with water, and 1 µL of the diluted PCR products was employed for the second sequential in-out PCR amplification reactions (**Table S3**). The resulting PCR product was visualized using a 1% agarose gel. Unpurified PCR products were forwarded for Sanger sequencing by Genewiz.

### Flow cytometry

J3-edited B cells in 5 mL FACS tubes (VWR) were identified by incubating with 1 µg of His-tagged HIV-1 JR-CSF gp120 at room temperature for 15 min. Then, cells were stained with appropriate antibodies, listed in **Table S4**, for 15 min in the dark at room temperature. If no intracellular staining was required, cells were then fixed with 4% paraformaldehyde (Electron Microscopy, Hatfield, PA). For intracellular staining, Cytofix/Cytoperm Fixation/Permeabilization Kit (BD Biosciences, Franklin Lakes, NJ) was used per manufacturer’s instructions. After staining, data were acquired with a FACSAria II (BD Biosciences) or Attune NxT Flow Cytometer (Thermo Fisher) and analyzed using FlowJo software version v10.7.1 (BD Biosciences).

### HCAb production

HCAb expression plasmids for J3-IgG1 and eCD4-IgG1 were designed to match the expected protein products after *IGHG* engineering (**Table S5**). J3-IgG1 was expressed from a PGK promoter in the plasmid pVax1 backbone (Thermo Fisher). eCD4-IgG1 was expressed from plasmid CMVR-eCD4-IgG2-v26 by replacing the IgG2 Fc region with human IgG1 Hi-CH2-CH3 sequences. Recombinant HCAbs were produced by transiently transfecting 293T cells as previously described^5^. In brief, 2.5×10^5^ 293T cells were seeded into one well of a 12-well plate and 1.5 μg of HCAb expression plasmids were transfected into cells using calcium phosphate transfection. For eCD4-IgG1 production, 0.4 μg of a human tyrosine protein sulfotranserase 2 (TPST2) expressing plasmid was co-transfected. The TPST2 expression plasmid was a gift from Michael Farzan (Harvard University).

### ELISAs

An ELISA protocol to detect total IgG (both HCAbs and natural IgG antibodies) was previously described^5^. In brief, polyclonal goat anti-human IgG antibody (Southern Biotech, Birmingham, AL) was used to coat plates at a 1:400 dilution, with anti-human IgG Fc HRP (Southern Biotech) at a 1:2000 dilution as a secondary antibody. Recombinant IgG1, Kappa from human myeloma (Millipore-Sigma), was serially diluted to make a standard curve in the range of 2000 – 15.625 ng/mL. HRP activity was measured by addition of SigmaFast OPD (Millipore-Sigma) and absorbance was measured at 450nm by a Mithras LB 940 plate reader (Berthold Technologies, Bad Wildbad, Germany).

The total IgG ELISA was modified to be specific for gp120 binding IgG and HCAbs by coating plates instead with 5 µg/mL recombinant HIV-1 JR-CSF gp120 protein (Immune Technology). Serially diluted recombinant J3-IgG1, produced from transfected 293T cells, was used to generate a standard curve from 400-3.125 ng/mL. This gp120/IgG ELISA was further modified to be specific for gp120/Igκ or gp120/Igλ by using either mouse anti-human Igκ HRP (Southern Biotech) or mouse anti-human Igλ HRP (Southern Biotech) as secondary antibodies at a 1:2000 dilution. Standard curves were prepared with recombinant heterodimers, in the range from 2000-31.25 ng/mL. For the gp120/Igκ ELISA, the recombinant heterodimer comprised J3-IgG1 and an Igκ Light chain antibody, Ofatumumab^5^, with knob-in-hole mutations^52^ included to force heterodimer formation ie S354C/T366W in the CH3 domain of J3-IgG1 and Y349C/T366S/L368A/Y407V in the CH3 domain of Ofatumumab. For the gp120/Igλ ELISA, recombinant heterodimer was produced in the same way between J3-IgG1 and an Igλ light chain antibody, Avelumab^53^.

### HIV-1 neutralization assay

HIV-1 pseudovirus strain BJOX2000.03.2 was produced from 293T cells as previously described^54^. Supernatant of edited Raji cells was collected and total IgG ELISA was used to quantify the J3-HCAb concentration in the supernatant. HIV neutralization activity of the J3-HCAb was measured by TZM-bl assay as previously described^55^. In brief, serially diluted J3-HCAb was mixed with pseudovirus. The input amount of pseudovirus was titrated to generate 10-20 times higher luminescence in wells with pseudovirus compared with background wells, containing no pseudovirus. Britelite Plus kit (Perkin-Elmer, Waltham, MA) was used to detect Luminescence which is measured by a Mithras LB 940 plate reader (Berthold Technologies).

### ADCC assay

An ADCC assay protocol was adapted from previous publications^25,56,57^. In brief, CEM.NKR.CCR5 cells^58^ were infected with HIV-1 strain NL4-3 to yield around 40-50% infected cells after 48 hours, by intracellular p24 staining, as described below. These target cells were stained with CellTrace Violet (Thermo Fisher) and resuspended in cRPMI at 2×10^6^ cells/mL. Then, 50 µL of target cells in U-bottom 96-well plates were incubated with 50 µL indicated HCAb or control antibodies (3BNC117 or PGT121, both from NIH HIV Reagent Program) (0.9 µg/ mL) for 15 min. NK-92 cells were resuspended in cRPMI at 2×10^6^ cells/mL and 100 µL added into each well (NK-92: target cell ratio of 2:1). The 96-well plates were spun at 300 x *g* for 1 min to encourage cell contact and incubated at 37°C for 4 hours. After incubation, cells were transferred into 5 mL FACS tubes, washed with PBS, stained with anti-CD56 antibody (BioLegend) and Near-IR fixable viability dye (Thermo Fisher) and incubated at room temperature for 15 min. Cells were washed and stained for intracellular p24 using p24/gag antibody KC57 (Beckman Coulter, Brea, CA) with the Cytofix/Cytoperm kit, as described above. Data were acquired on Attune NxT Flow Cytometer and analyzed by FlowJo.

### FcRn binding assay

HCAb-containing supernatants from edited Raji cells or transfected 293T cells were desalted and buffer-exchanged to PBS with Zeba™ Spin Desalting Columns, 40K MWCO, 5 mL (Thermo Fisher). The Lumit FcRn Binding Immunoassay kit (Promega, Madison, WI) was used to examine FcRn binding affinity according to the manufacturer’s instructions. In brief, desalted J3-HCAb protein was diluted with PBS and adjusted to pH 6. Control IgG-LgBit and hFcRn-smBiT were incubated with or without diluted J3-HCAb in 96-well half area plates (Corning, Corning, NY) at room temperature for 1 hour. Detection reagent was added to each well and incubated for 5 min. Luminescence was measured on a Mithras LB 940 plate reader (Berthold Technologies). During the assay, J3-HCAb competes with control IgG-LgBit to bind with hFcRn-smBiT and therefore decreases the luminescence signal. The following equation was used to calculate the percentage FcRn competition:

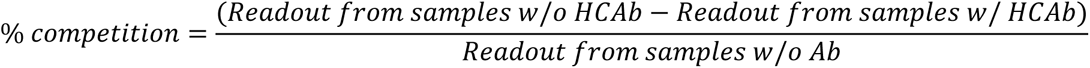

### Reverse transcription and digital droplet PCR (ddPCR)

mRNA was purified from edited or control B cells with RNeasy Plus Mini Kit (Qiagen) and used for reverse transcription to generate first-strand cDNA by SuperScript™ IV First-Strand Synthesis System (Thermo Fisher) with oligo d(T)_20_ primer per manufacturer’s instructions. The cDNA was mixed with ddPCR Supermix (Bio-Rad) and primer/probe sets to detect either total HCAbs or just the BCR or secreted isoforms (**Table S6**). The ddPCR mixture was formed into droplets using ddPCR droplet oil (Bio-Rad) and the PCR run on a C1000 Touch Thermal Cycler (Bio-Rad) with following conditions: 95°C 10 min, 40 cycles (94°C 1 min, 59°C 30 sec, 72°C 1 min), 98°C 10 min, and 4°C forever. The PCR products were read on a QX200 Droplet Reader (Bio-Rad) and analyzed with QuantaSoft analysis software (Bio-Rad).

### Statistics

Statistical analysis was performed by Prism v9.0.0 (GraphPad, La Jolla, CA), including paired or unpaired 2-tailed student’s t-test, simple linear regression with ANCOVA analysis, nonlinear least square regression and 1-way ANOVA with multiple comparisons. Hypothesis tests were 2-sided. The threshold of significance was set to 0.05.

### Illustrations

Figures 1a, 1c, 4a, 4c, 5a, 6a and Extended Figure 1c, 4d, 5a were created using BioRender.com.

## Acknowledgments

This research was supported by National Institutes of Health grants U19 HL156247, UM1 AI164561 and R01 AI167003 to P.M.C, and F30 AI186662 to A.M. Tonsil tissue was collected and processed by the Norris Comprehensive Cancer Center Translational Pathology Core, supported by National Cancer Institute grant P30 CA014089. Intellectual property discussed in this paper has been licensed by Be Biopharma from the University of Southern California. P.M.C is a paid member of Be Biopharma’s Scientific Advisory Board and has options to purchase stock in Be Biopharma.

---

**Extended Figure 1.**
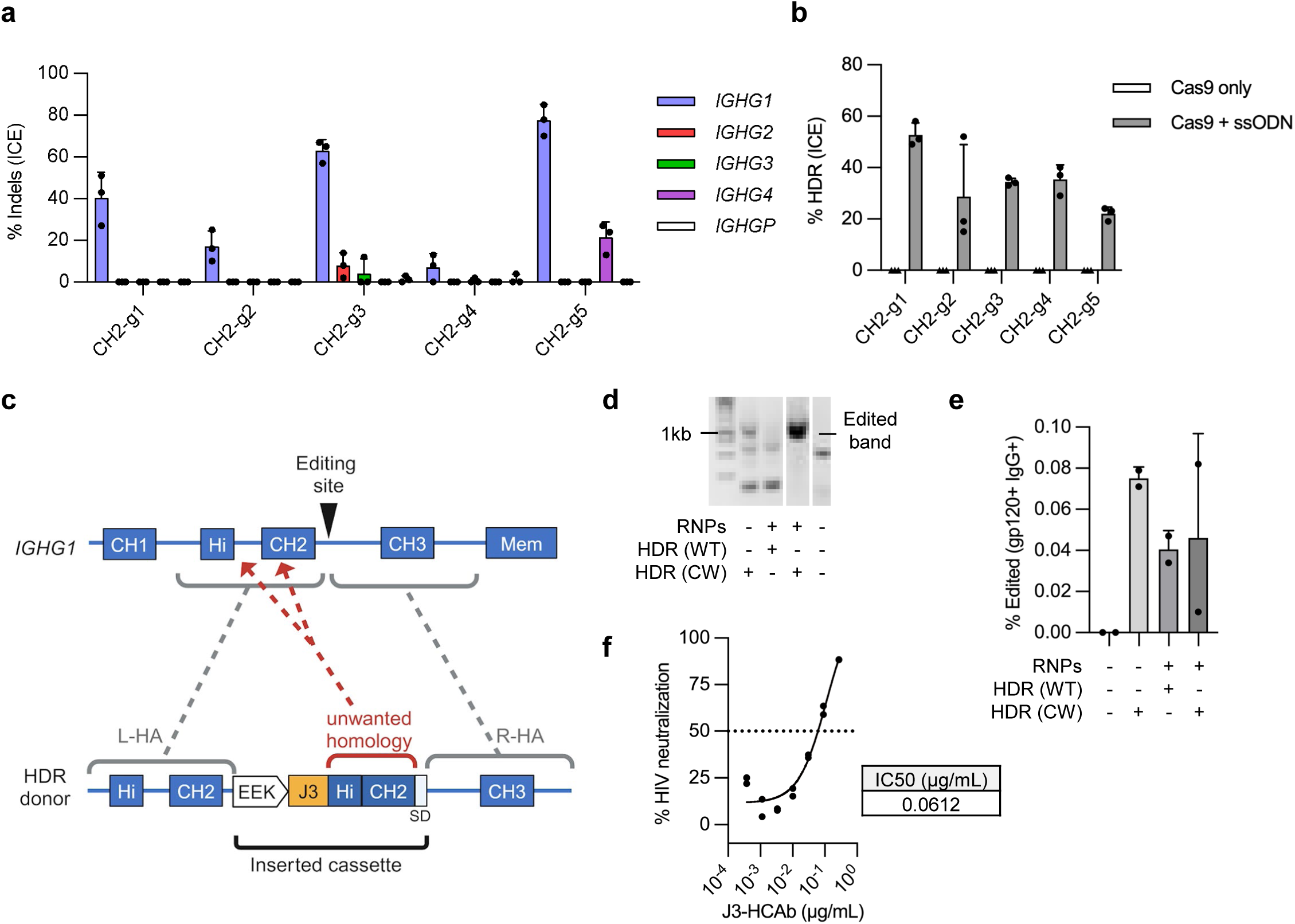
Development of editing approach for the *IGHG1* CH2-intron. (**a**) Five gRNAs targeting the CH2 intron (**Table S1**) were complexed with Cas9 protein and the RNPs electroporated into Raji cells. Indels generated at the on-target cut sites in *IGHG1* or potential off-target sites in other *IGHG* loci (*IGHG2* to P) were measured by Sanger sequencing and ICE analysis (*n* = 3). (**b**) Raji cells were edited with the panel of 5 CH2-intron Cas9 RNPs plus matched ssODN HDR templates carrying a 6-bp insert (**Table S1**). HDR efficiency was measured by Sanger sequencing and ICE analysis (*n* = 3). (**c**) Schematic showing how unmodified Hinge and CH2 sequences in the HDR template could interfere with the desired HDR editing outcome. The left-(L-HA) and right (R-HA) homology arms of the HDR template contain the genomic sequences flanking the Cas9 target site. (**d**) Raji cells were edited with CH2-g1 Cas9 RNPs plus plasmid HDR donors containing a B cell specific promoter, EEK, a J3 VHH as an ARD and either wild-type (WT) or codon-wobbled (CW) Hinge and CH2 exons. Efficient editing was detected by a specific nested in-out PCR at 7 days post-editing only when the CW HDR template was used. (**e**) Control plus edited Raji cells from (d) were analyzed by flow cytometry to detect cell surface J3-CH2-HCAb BCR. (**f**) J3-CH2-HCAb edited Raji cells were FACS-sorted for HIV-gp120 and IgG double positive cells and supernatants used in an HIV neutralization assay. Neutralization curves are shown for serial dilutions of supernatants and IC50 was calculated based on the curve (*n* = 2).

**Extended Figure 2.**
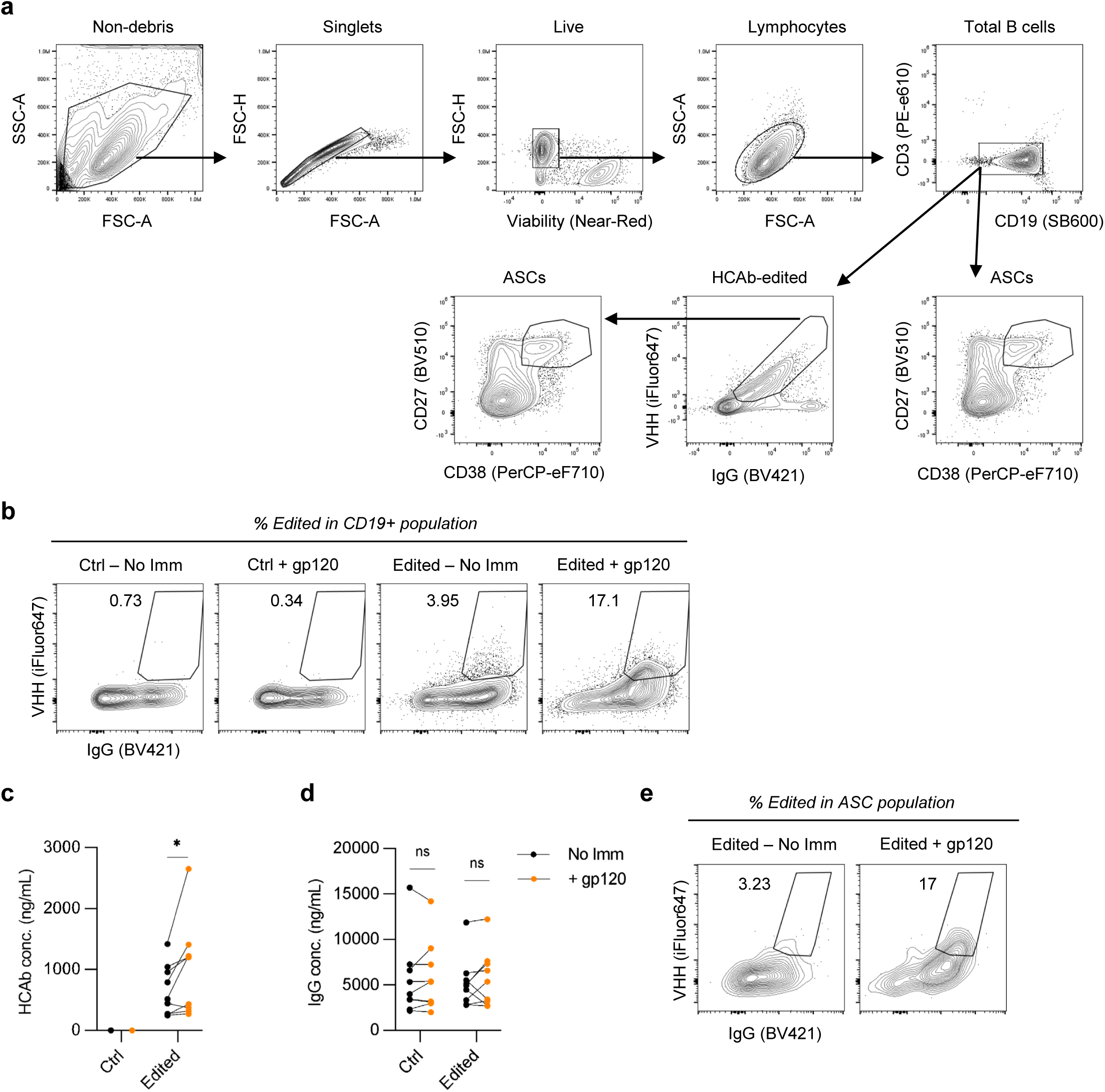
Analysis of J3-CH2 HCAb edited primary human B cells. (**a**) Gating strategy for flow cytometry analysis of J3-CH2 HCAb edited primary human B cells. Lymphocytes were identified by excluding debris, non-singlets, dead cells, and those of incorrect size. B cells were identified within lymphocytes as CD19+. The edited population was identified as VHH+ IgG+. ASCs in the CD19+ population were identified as CD27+CD38+. (**b**) Representative flow cytometry analysis of J3-CH2-HCAb edited B cells in tonsil organoids, showing increase in edited cells (VHH+ IgG+) within the total CD19+ B cell population after immunization; relates to **Fig 2g**. (**c**) J3-CH2 HCAbs and (**d**) total IgG in supernatant from matched tonsil organoid cultures without (No Imm) and with immunization (+ gp120) at day 12; relates to **Fig 2h**. (**e**) Representative flow cytometry, showing increase in edited population within the ASC population after immunization; relates to **Fig 2j**. Statistical analysis in (c, d) was paired 2-tailed t test between non-immunized and immunized samples. * *p* < 0.05, ns not significant.

**Extended figure 3.**
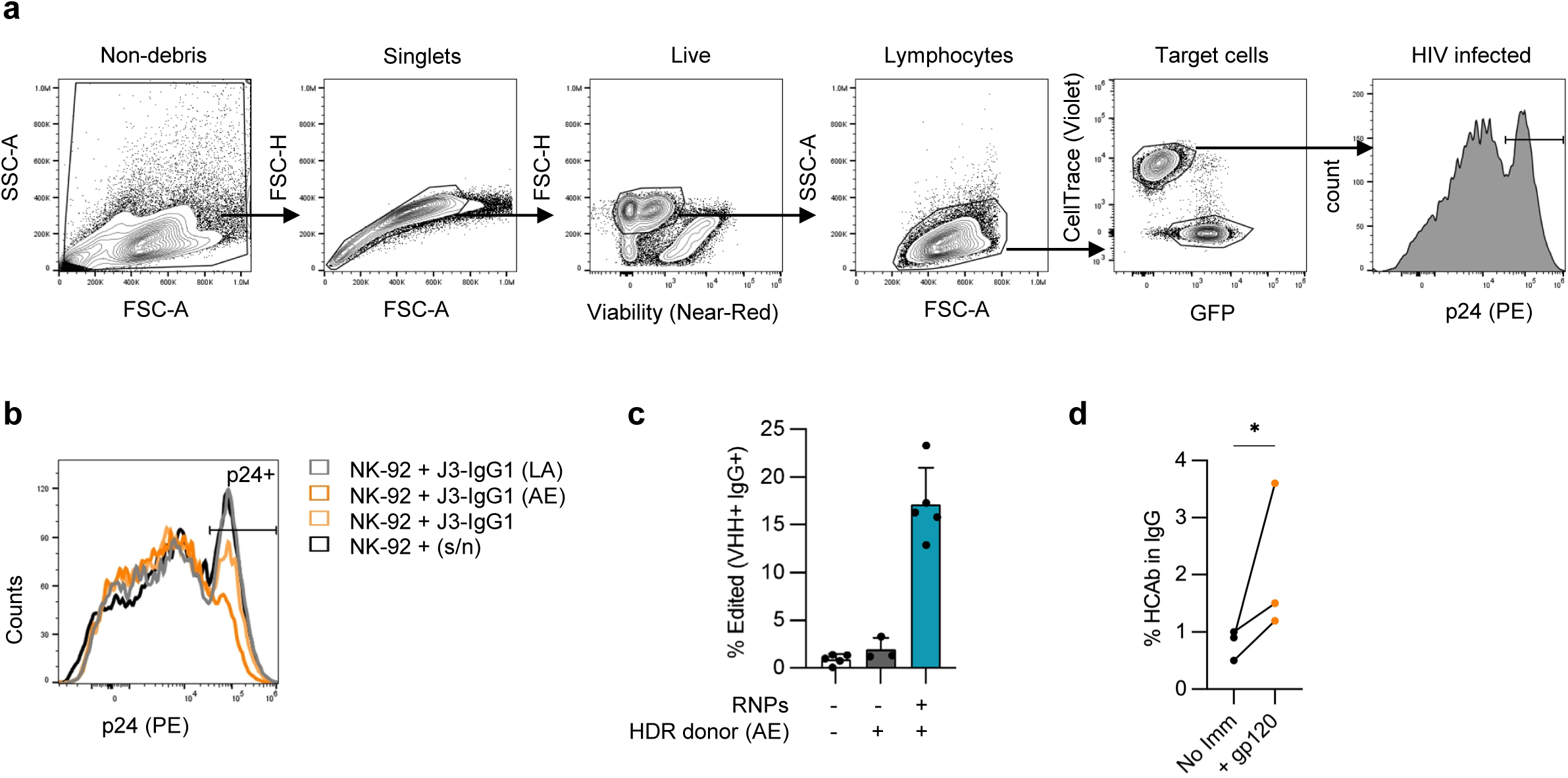
Analysis of CH2-HCAb edited cells and ADCC assays. (**a**) Representative gating scheme for ADCC assay. Live cells were identified by non-debris, singlets, viability dye and size. HIV-infected CEM.NKR.CCR5 target cells were stained with CellTrace and separated from GFP+ NK-92 effector cells. HIV infection rates in target cells were quantified by HIV p24 signal. (**b**) Representative graph showing changes in the percentage of infected (p24+) cells in the viable target cell population following incubation with NK-92 cells and the indicated Raji cell culture supernatants; relates to **Fig 3b**. (**c**) Editing efficiency of tonsil B cells with AE donor before organoid establishment. (**d**) Percentage of J3-CH2(AE) HCAbs in total IgG was measured in supernatants of tonsil organoid cultures containing AE-edited cells at day 12, with and without immunization, n=3 independent donors. Error bars represent mean ± SEM. Statistical analysis in (d) was a paired one-tailed t test. * *p* < 0.05.

**Extended figure 4.**
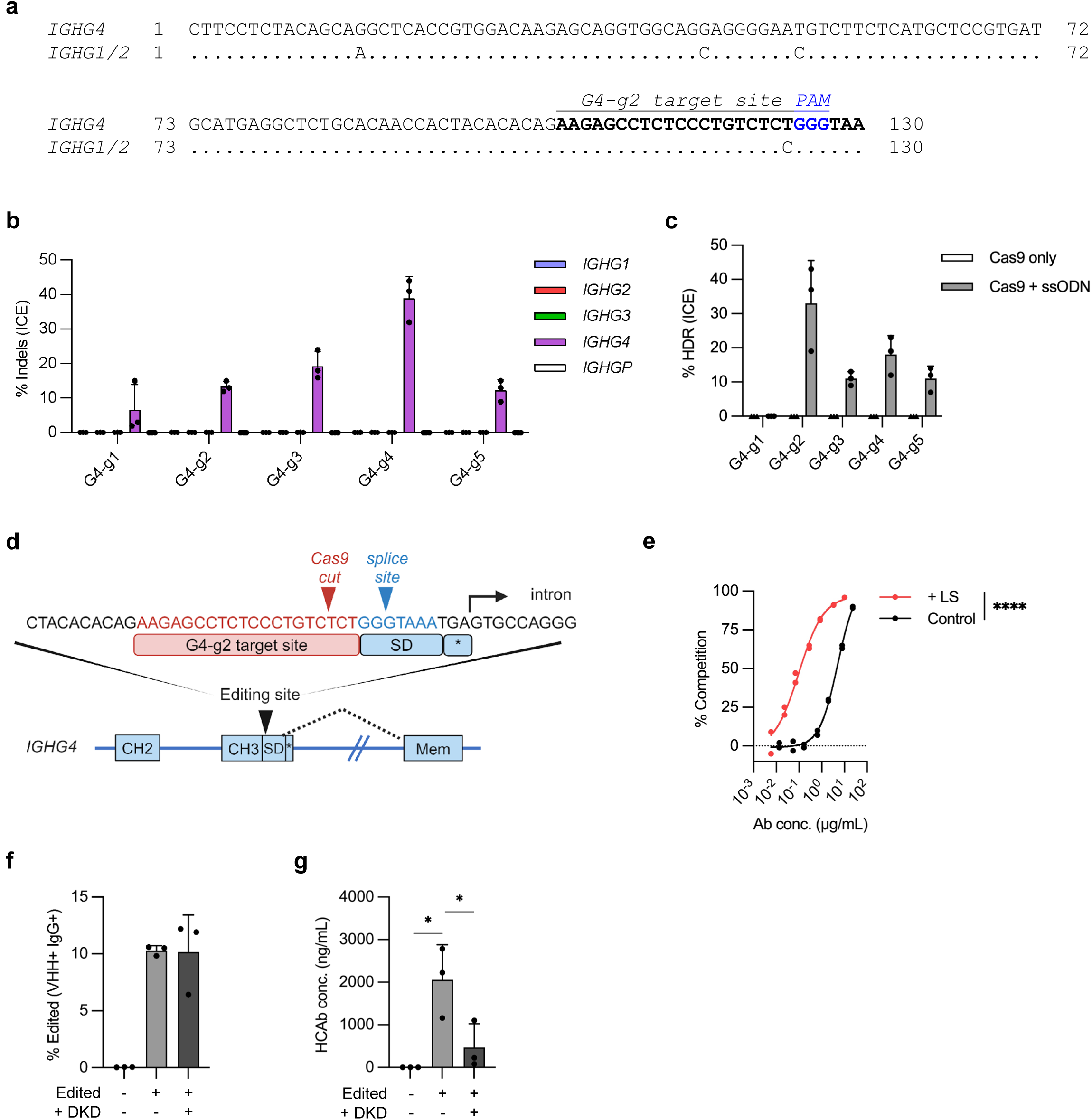
Development of editing approach targeting the *IGHG4* CH3 exon and FcRn assays. (**a**) Alignment of the *IGHG4* CH3 exon, upstream of the CH3 splice donor site, with the corresponding sequences in *IGHG1* and *IGHG2.* (**b**) Activity of 5 gRNAs targeting the *IGHG4* CH3-exon (Table S1) in Raji cells, with indels generated at the on-target sites in *IGHG4 or* potential off-target sites in other *IGHG* loci measured at day 7 post-editing by Sanger sequencing and ICE assay (*n* = 3). (**c**) Raji cells were edited with indicated Cas9 RNPs plus matched ssODN HDR templates carrying a 6-bp insert. HDR efficiency was measured by Sanger sequencing and ICE assay (*n* = 3) at day 7 post-editing. (**d**) Schematic showing the genomic sequence around the G4-g2 target site, 5 bp upstream from the CH3 SD. Stop codon is (*). (**e**) FcRn binding assay using J3-IgG1-HCAbs, with or without LS mutations, produced from transfected 293T cells. FcRn binding affinity was determined by competition with a control antibody and percentage competition calculated (*n* = 2). (**f-g**) Tonsil B cells were edited with G4-g2 Cas9 RNPs plus AAV6 HDR templates carrying J3-IgG1-HCAb expression cassette, with or without DKD mutations (n=3). Cells and supernatants were harvested at 2 days post-editing to measure (**f**) BCR expression by flow cytometry and (**g**) J3-IgG1-HCAb levels in supernatants by gp120-IgG ELISA. Error bars represent mean ± SEM. Statistical analysis were performed by nonlinear least square regression in (e) and unpaired 2-tailed t test in (f, g). * *p* < 0.05, ** *p* < 0.01, *** *p*<0.001, **** *p*<0.0001.

**Extended Figure 5.**
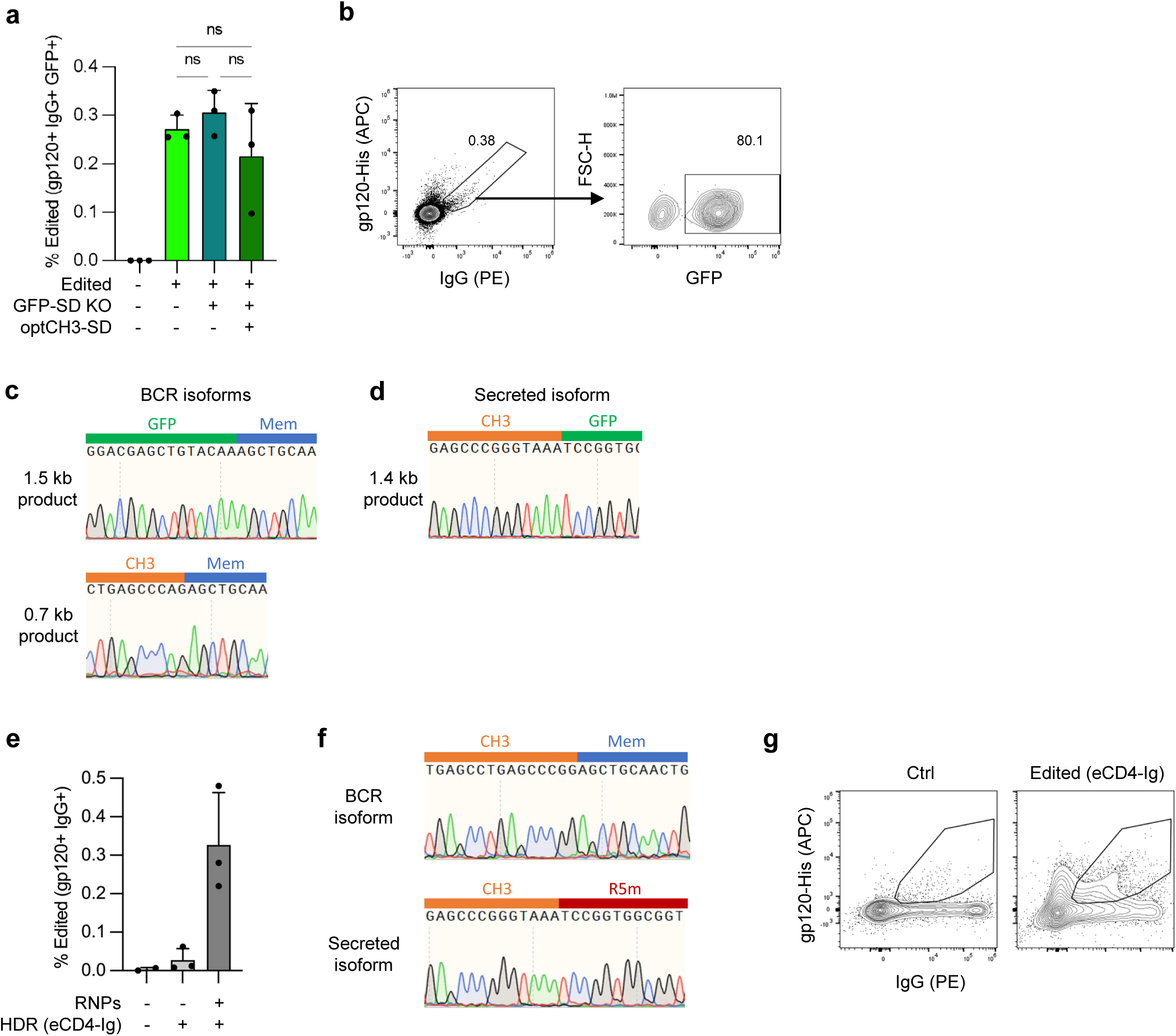
Incorporating additional domains at the C-terminus of HCAbs. (**a**) Raji cells were edited with G4-g2 Cas9 RNPs and plasmid HDR templates carrying the J3-IgG1-GFP cassette, with or without the GFP-SD KO and optimized CH3-SD mutations. Editing efficiency, defined as gp120+ IgG+ GFP+, was assessed by flow cytometry at day 7 post-electroporation (*n* = 3). (**b**) Gating strategy used to define edited cells. (**c**) Sanger sequencing to characterize BCR isoform RT-PCR products from **Fig 5c**. (**d**) Sanger sequencing to characterize secreted isoform RT-PCR products from **Fig 5d**. (**e**) Raji cells were electroporated with G4-g2 Cas9 RNPs plus plasmid HDR template for the eCD4-Ig cassette. Editing efficiency, defined as gp120+ IgG+, was detected at 7 days post-electroporation (*n* = 3). (**f**) Sanger sequencing to characterize BCR and secreted isoform RT-PCR products from **Fig 6b**, showing that the CCR5 peptide (R5m) was only present in the secreted form. (**g**) Representative gating strategy for detecting eCD4-Ig edited PBMC B cells, defined as gp120+ IgG+ in the CD19+ population; relates to **Fig 6c**. Error bars represent mean ± SEM. Statistical analysis was performed by 1-way ANOVA with multiple comparisons in (a). ns not significant.

