## Supplementary Tables for "Engineering B cells to Express Fully Customizable Antibodies with Enhanced Fc Functions"

Supplementary Table 1. gRNA sequences and corresponding ssODN HDR donors.
Supplementary Table 2. HDR donors to insert HCAbs at *IGHG1* CH2 intron or *IGHG4* CH3 exon.
Supplementary Table 3. PCR primer sets.
Supplementary Table 4. Flow cytometry staining panels.
Supplementary Table 5. Sequences of HCAb expression plasmids.
Supplementary Table 6. Primers and probes used for ddPCR.

**Supplementary Table 1. gRNA sequences and corresponding ssODN HDR donors.**

**a.** gRNAs targeting *IGHG1* CH2 intron

| gRNA | Sequence **PAM** | Matched ssODN HDR template (lowercase is XhoI restriction site) |
|---|---|---|
| CH2-g1 | ATGTGGCCCTCG CACCCCAC **GGG** | ATCGAGAAAACCATCTCCAAAGCCAAAGGTGGGACCCGTGctcgagGGGTGCGAG GGCCACATGGACAGAGGCCGGCTCGGCCCAC |
| CH2-g2 | AAGCCAAAGGTG GGACCCGT **GGG** | GCCGAGCCGGCCTCTGTCCATGTGGCCCTCGCACCCCACGctcgagGGTCCCAC CTTTGGCTTTGGAGATGGTTTTCTCGATGGGG |
| CH2-g3 | AGCCAAAGGTGG GACCCGTG **GGG** | GGCCGAGCCGGCCTCTGTCCATGTGGCCCTCGCACCCCACctcgagGGGTCCCA CCTTTGGCTTTGGAGATGGTTTTCTCGATGGG |
| CH2-g4 | GTGGGACCCGTG GGGTGCGA **GGG** | AGAGGGTGGGCCGAGCCGGCCTCTGTCCATGTGGCCCTCGctcgagCACCCCAC GGGTCCCACCTTTGGCTTTGGAGATGGTTTTC |
| CH2-g5 | CATGTGGCCCTC GCACCCCA **CGG** | TCGAGAAAACCATCTCCAAAGCCAAAGGTGGGACCCGTGGctcgagGGTGCGAGG GCCACATGGACAGAGGCCGGCTCGGCCCACC |

**b.** gRNAs targeting *IGHG4* CH3 exon

| gRNA | Sequence **PAM** | Matched ssODN HDR template (lowercase is XhoI restriction site) |
|---|---|---|
| G4-g1 | TCTCTGGGTAAAT GAGTGCC **AGG** | ACCCCGAGAGCCCGGGGAGCGGGGGCTTGCCGGCCCTGGCctcgagACTCATTT ACCCAGAGACAGGGAGAGGCTCTTCTGTGTGT |
| G4-g2 | AAGAGCCTCTCCC TGTCTCT **GGG** | GGAGCGGGGGCTTGCCGGCCCTGGCACTCATTTACCCAGActcgagGACAGGGA GAGGCTCTTCTGTGTGTAGTGGTTGTGCAGAG |
| G4-g3 | CGTGGACAAGAG CAGGTGGC **AGG** | TCATGCATCACGGAGCATGAGAAGACATTCCCCTCCTGCCctcgagACCTGCTCTT GTCCACGGTGAGCCTGCTGTAGAGGAAGAA |
| G4-g4 | GACAAGAGCAGG TGGCAGGA **GGG** | AGCCTCATGCATCACGGAGCATGAGAAGACATTCCCCTCCctcgagTGCCACCTG CTCTTGTCCACGGTGAGCCTGCTGTAGAGGA |
| G4-g5 | GGACAAGAGCAG GTGGCAGG **AGG** | GCCTCATGCATCACGGAGCATGAGAAGACATTCCCCTCCTctcgagGCCACCTGCT CTTGTCCACGGTGAGCCTGCTGTAGAGGAA |

**Supplementary Table 2. HDR donors to insert HCAbs at *IGHG1* CH2 intron or *IGHG4* CH3 exon.** (**a-c**) HDR donor sequences to create J3-CH2 HCAbs at the CH2-g1 target site in *IGHG1*, containing codon-wobbled IgG1 Hinge (Hi) and CH2 exons and a modified CH2 splice donor (modSD; underlined, with specific base change in lower case), and with additional (**b**) AE or (**c**) LA mutations (bold, underlined). (**d-f**) HDR donor sequences to create J3-IgG1 HCAbs at the G4-g2 target site in *IGHG4*, containing codon-wobbled IgG1 Hinge, CH2 and CH3 exons, and with additional (**e**) LS mutations (bold, underlined) or (**f**) DKD mutations (bold, underlined). (**g**) HDR donor for insertion of GFP domain at the C-terminus of CH3, using the G4-g2 gRNA target site in *IGHG4,* and inserting a J3-IgG1-GFP cassette containing a modified CH3-SD (modSD; underlined, with specific base change in lower case), an SGGG linker between CH3 and the GFP domain (italicized, underlined) and a modified GFP sequence (mGFP) with a cryptic SD at the 3' end removed (specific base change in lower case). (**h**) HDR for insertion of eCD4-IgG1 at the G4-g2 gRNA target site in *IGHG4.* The insert contains CD4 domains 1 and 3 as an ARD and a C-terminal CCR5 mimetic peptide (R5m), both derived from plasmid CMVR-eCD4-IgG2-v26, and an SGGG linker (italicized, underlined) between the CH3 and R5m sequences. Sequences are colored throughout to indicate specific regions, as noted in construct names, with multiple cloning sites between segments in black text. Additional abbreviations: LHA, left homology arm; RHA, right homology arm; EEK, EEK promoter; J3, anti-HIV VHH domain.

**a.** LHA-EEK-J3-HiCH2-modSD-**RHA**

ACTCATGCTCAGGGAGAGGGTCTTCTGGCTTTTTCCCAGGCTCTGGGCAGGCACAGGCTAGGTGCCCCTAACCCA
GGCCCTGCACACAAAGGGGCAGGTGCTGGGCTCAGACCTGCCAAGAGCCATATCCGGGAGGACCCTGCCCCTGAC
CTAAGCCCACCCCAAAGGCCAAACTCTCCACTCCCTCAGCTCGGACACCTTCTCTCCTCCCAGATTCCAGTAACTCC
CAATCTTCTCTCTGCAGAGCCCAAATCTTGTGACAAAACTCACACATGCCCACCGTGCCCAGGTAAGCCAGCCCAGG
CCTCGCCCTCCAGCTCAAGGCGGGACAGGTGCCCTAGAGTAGCCTGCATCCAGGGACAGGCCCCAGCCGGGTGCT
GACACGTCCACCTCCATCTCTTCCTCAGCACCTGAACTCCTGGGGGGACCGTCAGTCTTCCTCTTCCCCCCAAAACC
CAAGGACACCCTCATGATCTCCCGGACCCCTGAGGTCACATGCGTGGTGGTGGACGTGAGCCACGAAGACCCTGAG
GTCAAGTTCAACTGGTACGTGGACGGCGTGGAGGTGCATAATGCCAAGACAAAGCCGCGGGAGGAGCAGTACAACA
GCACGTACCGTGTGGTCAGCGTCCTCACCGTCCTGCACCAGGACTGGCTGAATGGCAAGGAGTACAAGTGCAAGGT
CTCCAACAAAGCCCTCCCAGCCCCCATCGAGAAAACCATCTCCAAAGCCAAAGGTGGGACCCGTGATCGATTAAACC
GGTGAGTTTCATGGTTACTTGCCTGAGAAGATTAAAAAAAGTAATGCTACCTTATGAGGGAGAGTCCCAGGGACCAA
GATAGCAACTGTCATAGCAACCGTCACACTGCTTTGGTCAAGGAGAAGACCCTTTGGGGAACTGAAAACAGAACCTT
GAGCACATCTGTTGCTTTCGCTCCCATCCTCCTCCAACAGGGCTGGGTGGAGCACTCCACACCCTTTCACCGGTCGT
ACGGCTCAGCCAGAGTAAAAATCACACCCATGACCTGGCCACTGAGGGCTTGATCAATTCACTTTGAATTTGGCATTA
AATACCATTAAGGTATATTAACTGATTTTAAAATAAGATATATTCGTGACCATGTTTTTAACTTTCAAAAATGTAGCTGC
CAGTGTGTGATTTTATTTCAGTTGTACAAAATATCTAAACCTATAGCAATGTGATTAATAAAAACTTAAACATATTTTCCA
GTACCTTAATTCTGTGATAGGAAAATTTTAATCTGAGTATTTTAATTTCATAATCTCTAAAATAGTTTAATGATTTGTCAT
TGTGTTGCTGTCGTTTACCCCAGCTGATCTCAAAAGTGATATTTAAGGAGATTATTTTGGTCTGCAACAACTTGATAGG
GCTCAGCCTCTCCCACCCAACGGGTGGAATCCCCCAGAGGGGGATTTCCAAGAGGCCACCTGGCAGTTGCTGAGG
GTCAGAAGTGAAGCTAGCCACTTCCTCTTAGGCAGGTGGCCAAGATTACAGTTGACCCGTACGTGCAGCTGTGCCCA
GCCTGCCCCATCCCCTGCTCATTTGCATGTTCCCAGAGCACAACCTCCTGCCCTGAAGCCTTATTAATAGGCTGGTC
ACACTTTGTGCAGGAGTCAGACTCAGTCAGGACACAGCTGGATCCACTAGTCCAGTGTGGTGGAATTCACCATGGAG
TTTGGGCTGAGCTGGCTTTTTCTTGTGGCTATTTTAAAAGGTGTCCAGTGTGAGGTGCAGCTGGTGGAGTCTGGGGG
AGGCTTGGTACAGGCCGGGGGGTTCCTGAGACTCTCCTGTGAGCTGAGGGGAAGCATCTTTAACCAGTATGCCATG
GCCTGGTTCCGCCAGGCTCCAGGGAAGGAGAGGGAGTTCGTCGCCGGCATGGGCGCCGTGCCCCACTACGGCGA
GTTCGTGAAGGGCCGGTTCACCATCTCCAGAGACAATGCCAAGAGCACGGTGTATCTGCAAATGAGCAGCCTGAAG
CCCGAGGACACGGCCATCTATTTCTGTGCCAGGAGCAAGAGCACCTACATCAGCTACAACAGCAACGGCTACGACTA
CTGGGGCAGGGGAACCCAGGTCACCGTGAGCAGCGAGCCGAAGAGCTGTGACAAAACACATACATGCCCCCCCTG
CCCCGCGCCCGAGCTGCTTGGCGGTCCAAGCGTGTTCCTGTTCCCCCCCAAGCCCAAGGACACCCTGATGATCAGC
AGAACCCCCGAGGTGACCTGCGTGGTGGTGGACGTGAGCCACGAGGACCCCGAGGTGAAGTTCAACTGGTACGTG
GACGGCGTGGAGGTGCACAACGCCAAGACCAAGCCTAGAGAGGAGCAGTACAACAGCACCTACAGAGTGGTGAGC
GTGCTGACCGTGCTGCACCAAGACTGGCTGAACGGCAAGGAGTACAAGTGCAAGGTGTCCAACAAGGCCCTGCCCG
CCCCCATCGAGAAGACCATCAGCAAGGCCAAAGGTGaGAGCCGTGGAGCTC**GGGTGCGAGGGCCACATGGACAGA**

**GGCCGGCTCGGCCCACCCTCTGCCCTGAGAGTGACCGCTGTACCAACCTCTGTCCCTACAGGGCAGCCCCGAGA**
**ACCACAGGTGTACACCCTGCCCCCATCCCGGGATGAGCTGACCAAGAACCAGGTCAGCCTGACCTGCCTGGTCAA**
**AGGCTTCTATCCCAGCGACATCGCCGTGGAGTGGGAGAGCAATGGGCAGCCGGAGAACAACTACAAGACCACGC**
**CTCCCGTGCTGGACTCCGACGGCTCCTTCTTCCTCTACAGCAAGCTCACCGTGGACAAGAGCAGGTGGCAGCAGG**
**GGAACGTCTTCTCATGCTCCGTGATGCATGAGGCTCTGCACAACCACTACACACAGAAGAGCCTCTCCCTGTCTCC**
**GGGTAAATGAGTGCCACGGCCGGCAAGCCCCCGCTCCCCAGGCTCTCGGGGTCGCGCGAGGATGCTTGGCACGT**
**ACCCCGTGTACATACTTCCCAGGCACCCAGCATGGAAATAAAGCACCCAGCGCTTCCCTGGGCCCCTGCGAGACT**
**GTGATGGTTCTTTCCACGGGTCAGGCCGAGTCTGAGGCCTGAGTGGCATGAGGGAGGCAGAGTGGGTCCCACTGT**
**CCCCACACTGGCCCAGGCTGTGCAGGTGTGCCTGGGCCGCCTAGGGTGGGGCTCAGCCAGGGGCTGCCCTCGGC**
**AGGGTGGGGGATTTGCCAGCGTGGCCCTCCCTCCAGCAGCAGCTGCCCTGGGCT**

**b.** LHA-EEK-J3-HiCH2(**AE**)-modSD-**RHA**

ACTCATGCTCAGGGAGAGGGTCTTCTGGCTTTTTCCCCAGGCTCTGGGCAGGCACAGGCTAGGTGCCCCTAACCCA
GGCCCTGCACACAAAGGGGCAGGTGCTGGGCTCAGACCTGCCAAGAGCCATATCCGGGAGGACCCTGCCCCTGAC
CTAAGCCCACCCCAAAGGCCAAACTCTCCACTCCCTCAGCTCGGACACCTTCTCTCCTCCCAGATTCCAGTAACTCC
CAATCTTCTCTCTGCAGAGCCCAAATCTTGTGACAAAACTCACACATGCCCACCGTGCCCAGGTAAGCCAGCCCAGG
CCTCGCCCTCCAGCTCAAGGCGGGACAGGTGCCCTAGAGTAGCCTGCATCCAGGGACAGGCCCCAGCCGGGTGCT
GACACGTCCACCTCCATCTCTTCCTCAGCACCTGAACTCCTGGGGGGACCGTCAGTCTTCCTCTTCCCCCCAAAACC
CAAGGACACCCTCATGATCTCCCGGACCCCTGAGGTCACATGCGTGGTGGTGGACGTGAGCCACGAAGACCCTGAG
GTCAAGTTCAACTGGTACGTGGACGGCGTGGAGGTGCATAATGCCAAGACAAAGCCGCGGGAGGAGCAGTACAACA
GCACGTACCGTGTGGTCAGCGTCCTCACCGTCCTGCACCAGGACTGGCTGAATGGCAAGGAGTACAAGTGCAAGGT
CTCCAACAAAGCCCTCCCAGCCCCCATCGAGAAAACCATCTCCAAAGCCAAAGGTGGGACCCGTGATCGATTAAACC
GGTGAGTTTCATGGTTACTTGCCTGAGAAGATTAAAAAAAGTAATGCTACCTTATGAGGGAGAGTCCCAGGGACCAA
GATAGCAACTGTCATAGCAACCGTCACACTGCTTTGGTCAAGGAGAAGACCCTTTGGGGAACTGAAAACAGAACCTT
GAGCACATCTGTTGCTTTCGCTCCCATCCTCCTCCAACAGGGCTGGGTGGAGCACTCCACACCCTTTCACCGGTCGT
ACGGCTCAGCCAGAGTAAAAATCACACCCATGACCTGGCCACTGAGGGCTTGATCAATTCACTTTGAATTTGGCATTA
AATACCATTAAGGTATATTAACTGATTTTAAAATAAGATATATTCGTGACCATGTTTTTAACTTTCAAAAATGTAGCTGC
CAGTGTGTGATTTTATTTCAGTTGTACAAAATATCTAAACCTATAGCAATGTGATTAATAAAAACTTAAACATATTTTCCA
GTACCTTAATTCTGTGATAGGAAAATTTTAATCTGAGTATTTTAATTTCATAATCTCTAAAATAGTTTAATGATTTGTCAT
TGTGTTGCTGTCGTTTACCCCAGCTGATCTCAAAAGTGATATTTAAGGAGATTATTTTGGTCTGCAACAACTTGATAGG
GCTCAGCCTCTCCCACCCAACGGGTGGAATCCCCCAGAGGGGGATTTCCAAGAGGCCACCTGGCAGTTGCTGAGG
GTCAGAAGTGAAGCTAGCCACTTCCTCTTAGGCAGGTGGCCAAGATTACAGTTGACCCGTACGTGCAGCTGTGCCCA
GCCTGCCCCATCCCCTGCTCATTTGCATGTTCCCAGAGCACAACCTCCTGCCCTGAAGCCTTATTAATAGGCTGGTC
ACACTTTGTGCAGGAGTCAGACTCAGTCAGGACACAGCTGGATCCACTAGTCCAGTGTGGTGGAATTCACCATGGAG
TTTGGGCTGAGCTGGCTTTTTCTTGTGGCTATTTTAAAAGGTGTCCAGTGTGAGGTGCAGCTGGTGGAGTCTGGGGG
AGGCTTGGTACAGGCCGGGGGGTTCCTGAGACTCTCCTGTGAGCTGAGGGGAAGCATCTTTAACCAGTATGCCATG
GCCTGGTTCCGCCAGGCTCCAGGGAAGGAGAGGGAGTTCGTCGCCGGCATGGGCGCCGTGCCCCACTACGGCGA
GTTCGTGAAGGGCCGGTTCACCATCTCCAGAGACAATGCCAAGAGCACGGTGTATCTGCAAATGAGCAGCCTGAAG
CCCGAGGACACGGCCATCTATTTCTGTGCCAGGAGCAAGAGCACCTACATCAGCTACAACAGCAACGGCTACGACTA
CTGGGGCAGGGGAACCCAGGTCACCGTGAGCAGCGAGCCGAAGAGCTGTGACAAAACACATACATGCCCCCCCTG
CCCCGCGCCCGAGCTGCTT**GCC**GGTCCC**GAC**GTGTTCCTGTTCCCCCCCAAGCCCAAGGACACCCTGATGATCAGC
AGAACCCCCGAGGTGACCTGCGTGGTGGTGGACGTGAGCCACGAGGACCCCGAGGTGAAGTTCAACTGGTACGTG
GACGGCGTGGAGGTGCACAACGCCAAGACCAAGCCTAGAGAGGAGCAGTACAACAGCACCTACAGAGTGGTGAGC
GTGCTGACCGTGCTGCACCAAGACTGGCTGAACGGCAAGGAGTACAAGTGCAAGGTGTCCAACAAGGCCCTGCCC**C**
**TG**CCC**GAG**GAGAAGACCATCAGCAAGGCCAAAGGTGaGAGCCGTGGAGCTC**GGGTGCGAGGGCCACATGGACAG**
**AGGCCGGCTCGGCCCACCCTCTGCCCTGAGAGTGACCGCTGTACCAACCTCTGTCCCTACAGGGCAGCCCCGAG**
**AACCACAGGTGTACACCCTGCCCCCATCCCGGGATGAGCTGACCAAGAACCAGGTCAGCCTGACCTGCCTGGTCA**
**AAGGCTTCTATCCCAGCGACATCGCCGTGGAGTGGGAGAGCAATGGGCAGCCGGAGAACAACTACAAGACCACG**
**CCTCCCGTGCTGGACTCCGACGGCTCCTTCTTCCTCTACAGCAAGCTCACCGTGGACAAGAGCAGGTGGCAGCAG**
**GGGAACGTCTTCTCATGCTCCGTGATGCATGAGGCTCTGCACAACCACTACACACAGAAGAGCCTCTCCCTGTCTC**
**CGGGTAAATGAGTGCCACGGCCGGCAAGCCCCCGCTCCCCAGGCTCTCGGGGTCGCGCGAGGATGCTTGGCACG**
**TACCCCGTGTACATACTTCCCAGGCACCCAGCATGGAAATAAAGCACCCAGCGCTTCCCTGGGCCCCTGCGAGAC**
**TGTGATGGTTCTTTCCACGGGTCAGGCCGAGTCTGAGGCCTGAGTGGCATGAGGGAGGCAGAGTGGGTCCCACTG**

**TCCCCACACTGGCCCAGGCTGTGCAGGTGTGCCTGGGCCGCCTAGGGTGGGGCTCAGCCAGGGGCTGCCCTCGG CAGGGTGGGGGATTTGCCAGCGTGGCCCTCCCTCCAGCAGCAGCTGCCCTGGGCT**

**c.** LHA-EEK-J3-HiCH2(**LA**)-modSD-**RHA**

ACTCATGCTCAGGGAGAGGGTCTTCTGGCTTTTTCCCCAGGCTCTGGGCAGGCACAGGCTAGGTGCCCCTAACCCA GGCCCTGCACACAAAGGGGCAGGTGCTGGGCTCAGACCTGCCAAGAGCCATATCCGGGAGGACCCTGCCCCTGAC CTAAGCCCACCCCAAAGGCCAAACTCTCCACTCCCTCAGCTCGGACACCTTCTCTCCTCCCAGATTCCAGTAACTCC CAATCTTCTCTCTGCAGAGCCCAAATCTTGTGACAAAACTCACACATGCCCACCGTGCCCAGGTAAGCCAGCCCAGG CCTCGCCCTCCAGCTCAAGGCGGGACAGGTGCCCTAGAGTAGCCTGCATCCAGGGACAGGCCCCAGCCGGGTGCT GACACGTCCACCTCCATCTCTTCCTCAGCACCTGAACTCCTGGGGGGACCGTCAGTCTTCCTCTTCCCCCCAAAACC CAAGGACACCCTCATGATCTCCCGGACCCCTGAGGTCACATGCGTGGTGGTGGACGTGAGCCACGAAGACCCTGAG GTCAAGTTCAACTGGTACGTGGACGGCGTGGAGGTGCATAATGCCAAGACAAAGCCGCGGGAGGAGCAGTACAACA GCACGTACCGTGTGGTCAGCGTCCTCACCGTCCTGCACCAGGACTGGCTGAATGGCAAGGAGTACAAGTGCAAGGT CTCCAACAAAGCCCTCCCAGCCCCCATCGAGAAAACCATCTCCAAAGCCAAAGGTGGGACCCGTGATCGATTAAACC GGTGAGTTTCATGGTTACTTGCCTGAGAAGATTAAAAAAAGTAATGCTACCTTATGAGGGAGAGTCCCAGGGACCAA GATAGCAACTGTCATAGCAACCGTCACACTGCTTTGGTCAAGGAGAAGACCCTTTGGGGAACTGAAAACAGAACCTT GAGCACATCTGTTGCTTTCGCTCCCATCCTCCTCCAACAGGGCTGGGTGGAGCACTCCACACCCTTTCACCGGTCGT ACGGCTCAGCCAGAGTAAAAATCACACCCATGACCTGGCCACTGAGGGCTTGATCAATTCACTTTGAATTTGGCATTA AATACCATTAAGGTATATTAACTGATTTTAAAATAAGATATATTCGTGACCATGTTTTTAACTTTCAAAAATGTAGCTGC CAGTGTGTGATTTTATTTCAGTTGTACAAAATATCTAAACCTATAGCAATGTGATTAATAAAAACTTAAACATATTTTCCA GTACCTTAATTCTGTGATAGGAAAATTTTAATCTGAGTATTTTAATTTCATAATCTCTAAAATAGTTTAATGATTTGTCAT TGTGTTGCTGTCGTTTACCCCAGCTGATCTCAAAAGTGATATTTAAGGAGATTATTTTGGTCTGCAACAACTTGATAGG GCTCAGCCTCTCCCACCCAACGGGTGGAATCCCCCAGAGGGGGATTTCCAAGAGGCCACCTGGCAGTTGCTGAGG GTCAGAAGTGAAGCTAGCCACTTCCTCTTAGGCAGGTGGCCAAGATTACAGTTGACCCGTACGTGCAGCTGTGCCCA GCCTGCCCCATCCCCTGCTCATTTGCATGTTCCCAGAGCACAACCTCCTGCCCTGAAGCCTTATTAATAGGCTGGTC ACACTTTGTGCAGGAGTCAGACTCAGTCAGGACACAGCTGGATCCACTAGTCCAGTGTGGTGGAATTCACCATGGAG TTTGGGCTGAGCTGGCTTTTTCTTGTGGCTATTTTAAAAGGTGTCCAGTGTGAGGTGCAGCTGGTGGAGTCTGGGGG AGGCTTGGTACAGGCCGGGGGGTTCCTGAGACTCTCCTGTGAGCTGAGGGGAAGCATCTTTAACCAGTATGCCATG GCCTGGTTCCGCCAGGCTCCAGGGAAGGAGAGGGAGTTCGTCGCCGGCATGGGCGCCGTGCCCCACTACGGCGA GTTCGTGAAGGGCCGGTTCACCATCTCCAGAGACAATGCCAAGAGCACGGTGTATCTGCAAATGAGCAGCCTGAAG CCCGAGGACACGGCCATCTATTTCTGTGCCAGGAGCAAGAGCACCTACATCAGCTACAACAGCAACGGCTACGACTA CTGGGGCAGGGGAACCCAGGTCACCGTGAGCAGCGAGCCGAAGAGCTGTGACAAAACACATACATGCCCCCCCTG CCCCGCGCCCGAG**GCTGCA**GGCGGTCCAAGCGTGTTCCTGTTCCCCCCCAAGCCCAAGGACACCCTGATGATCAG CAGAACCCCCGAGGTGACCTGCGTGGTGGTGGACGTGAGCCACGAGGACCCCGAGGTGAAGTTCAACTGGTACGT GGACGGCGTGGAGGTGCACAACGCCAAGACCAAGCCTAGAGAGGAGCAGTACAACAGCACCTACAGAGTGGTGAG CGTGCTGACCGTGCTGCACCAAGACTGGCTGAACGGCAAGGAGTACAAGTGCAAGGTGTCCAACAAGGCCCTGGGA GCCCCCATCGAGAAGACCATCAGCAAGGCCAAAGGTGaGAGCCGTGGAGCTC**GGGTGCGAGGGCCACATGGACAG AGGCCGGCTCGGCCCACCCTCTGCCCTGAGAGTGACCGCTGTACCAACCTCTGTCCCTACAGGGCAGCCCCGAG AACCACAGGTGTACACCCTGCCCCCATCCCGGGATGAGCTGACCAAGAACCAGGTCAGCCTGACCTGCCTGGTCA AAGGCTTCTATCCCAGCGACATCGCCGTGGAGTGGGAGAGCAATGGGCAGCCGGAGAACAACTACAAGACCACG CCTCCCGTGCTGGACTCCGACGGCTCCTTCTTCCTCTACAGCAAGCTCACCGTGGACAAGAGCAGGTGGCAGCAG GGGAACGTCTTCTCATGCTCCGTGATGCATGAGGCTCTGCACAACCACTACACACAGAAGAGCCTCTCCCTGTCTC CGGGTAAATGAGTGCCACGGCCGGCAAGCCCCCGCTCCCCAGGCTCTCGGGGTCGCGCGAGGATGCTTGGCACG TACCCCGTGTACATACTTCCCAGGCACCCAGCATGGAAATAAAGCACCCAGCGCTTCCCTGGGCCCCTGCGAGAC TGTGATGGTTCTTTCCACGGGTCAGGCCGAGTCTGAGGCCTGAGTGGCATGAGGGAGGCAGAGTGGGTCCCACTG TCCCCACACTGGCCCAGGCTGTGCAGGTGTGCCTGGGCCGCCTAGGGTGGGGCTCAGCCAGGGGCTGCCCTCGG CAGGGTGGGGGATTTGCCAGCGTGGCCCTCCCTCCAGCAGCAGCTGCCCTGGGCT**

**d.** LHA-EEK-J3-HiCH2CH3-**RHA**

ATCTCTTCCTCAGCACCTGAGTTCCTGGGGGGACCATCAGTCTTCCTGTTCCCCCCAAAACCCAAGGACACTCTCAT GATCTCCCGGACCCCTGAGGTCACGTGCGTGGTGGTGGACGTGAGCCAGGAAGACCCCGAGGTCCAGTTCAACTG GTACGTGGATGGCGTGGAGGTGCATAATGCCAAGACAAAGCCGCGGGAGGAGCAGTTCAACAGCACGTACCGTGT GGTCAGCGTCCTCACCGTCCTGCACCAGGACTGGCTGAACGGCAAGGAGTACAAGTGCAAGGTCTCCAACAAAGGC

CTCCCGTCCTCCATCGAGAAAACCATCTCCAAAGCCAAAGGTGGGACCCACGGGGTGCGAGGGCCACATGGACAGA
GGTCAGCTCGGCCCACCCTCTGCCCTGGGAGTGACCGCTGTGCCAACCTCTGTCCCTACAGGGCAGCCCCGAGAG
CCACAGGTGTACACCCTGCCCCCATCCCAGGAGGAGATGACCAAGAACCAGGTCAGCCTGACCTGCCTGGTCAAAG
GCTTCTACCCCAGCGACATCGCCGTGGAGTGGGAGAGCAATGGGCAGCCGGAGAACAACTACAAGACCACGCCTC
CCGTGCTGGACTCCGACGGCTCCTTCTTCCTCTACAGCAGGCTCACCGTGGACAAGAGCAGGTGGCAGGAGGGGA
ATGTCTTCTCATGCTCCGTGATGCATGAGGCTCTGCACAACCACTACACACAGAAGAGCCTCTCCCTGTCATCGATTA
AACCGGTGAGTTTCATGGTTACTTGCCTGAGAAGATTAAAAAAAGTAATGCTACCTTATGAGGGAGAGTCCCAGGGA
CCAAGATAGCAACTGTCATAGCAACCGTCACACTGCTTTGGTCAAGGAGAAGACCCTTTGGGGAACTGAAAACAGAA
CCTTGAGCACATCTGTTGCTTTCGCTCCCATCCTCCTCCAACAGGGCTGGGTGGAGCACTCCACACCCTTTCACCGG
TCGTACGGCTCAGCCAGAGTAAAAATCACACCCATGACCTGGCCACTGAGGGCTTGATCAATTCACTTTGAATTTGG
CATTAAATACCATTAAGGTATATTAACTGATTTTAAAATAAGATATATTCGTGACCATGTTTTTAACTTTCAAAAATGTAG
CTGCCAGTGTGTGATTTTATTTCAGTTGTACAAAATATCTAAACCTATAGCAATGTGATTAATAAAAACTTAAACATATT
TTCCAGTACCTTAATTCTGTGATAGGAAAATTTTAATCTGAGTATTTTAATTTCATAATCTCTAAAATAGTTTAATGATTT
GTCATTGTGTTGCTGTCGTTTACCCCAGCTGATCTCAAAAGTGATATTTAAGGAGATTATTTTGGTCTGCAACAACTTG
ATAGGGCTCAGCCTCTCCCACCCAACGGGTGGAATCCCCCAGAGGGGGATTTCCAAGAGGCCACCTGGCAGTTGCT
GAGGGTCAGAAGTGAAGCTAGCCACTTCCTCTTAGGCAGGTGGCCAAGATTACAGTTGACCCGTACGTGCAGCTGT
GCCCAGCCTGCCCCATCCCCTGCTCATTTGCATGTTCCCAGAGCACAACCTCCTGCCCTGAAGCCTTATTAATAGGC
TGGTCACACTTTGTGCAGGAGTCAGACTCAGTCAGGACACAGCTGGATCCACTAGTCCAGTGTGGTGGAATTCACCA
TGGAGTTTGGGCTGAGCTGGCTTTTTCTTGTGGCTATTTTAAAAGGTGTCCAGTGTGAGGTGCAGCTGGTGGAGTCT
GGGGGAGGCTTGGTACAGGCCGGGGGGTTCCTGAGACTCTCCTGTGAGCTGAGGGGAAGCATCTTTAACCAGTATG
CCATGGCCTGGTTCCGCCAGGCTCCAGGGAAGGAGAGGGAGTTCGTCGCCGGCATGGGCGCCGTGCCCCACTACG
GCGAGTTCGTGAAGGGCCGGTTCACCATCTCCAGAGACAATGCCAAGAGCACGGTGTATCTGCAAATGAGCAGCCT
GAAGCCCGAGGACACGGCCATCTATTTCTGTGCCAGGAGCAAGAGCACCTACATCAGCTACAACAGCAACGGCTAC
GACTACTGGGGCAGGGGAACCCAGGTCACCGTGAGCAGCGAGCCGAAGAGCTGTGACAAAACACATACATGCCCC
CCCTGCCCCGCGCCCGAGCTGCTTGGCGGTCCAAGCGTGTTCCTGTTCCCCCCCAAGCCCAAGGACACCCTGATGA
TCAGCAGAACCCCCGAGGTGACCTGCGTGGTGGTGGACGTGAGCCACGAGGACCCCGAGGTGAAGTTCAACTGGT
ACGTGGACGGCGTGGAGGTGCACAACGCCAAGACCAAGCCTAGAGAGGAGCAGTACAACAGCACCTACAGAGTGG
TGAGCGTGCTGACCGTGCTGCACCAAGACTGGCTGAACGGCAAGGAGTACAAGTGCAAGGTGTCCAACAAGGCCCT
GCCCGCCCCCATCGAGAAGACCATCAGCAAGGCCAAAGGGCAGCCTAGAGAGCCCCAAGTGTACACCCTGCCCCC
TAGCAGAGACGAGCTGACCAAGAACCAAGTGAGCCTGACCTGCCTGGTGAAGGGCTTCTACCCTAGCGACATCGCC
GTGGAGTGGGAGAGCAACGGGCAGCCCGAGAACAACTACAAGACCACCCCCCCCGTGCTGGACAGCGACGGCAGC
TTCTTCCTGTACAGCAAGCTGACCGTGGACAAGAGCAGATGGCAGCAAGGCAACGTGTTCAGCTGCAGCGTGATGC
ACGAGGCCCTGCACAACCACTACACACAGAAGAGCCTGAGCCTGAGC**CCGGGTAAATGAGTGCCAGGGCCGGCAA**
**GCCCCCGCTCCCCGGGCTCTCGGGGTCGCGCGAGGATGCTTGGCACGTACCCCGTGTACATACTTCCCGGGCGC**
**CCAGCATGGAAATAAAGCACCCAGCGCTGCCCTGGGCCCCTGCGAGACTGTGATGGTTCTTTCCACGGGTCAGGC**
**CGAGTCTGAGGCCTGAGTGGCATGAGGGAGGCAGAGCGGGTCCCACTGTCCCCACACTGGCCCAGGCTGTGCAG**
**GTGTGCCTGGGCCGCCTAGGGTGGGGCTCAGCCAGGGGCTGCCCTCGGCAGGGTGGGGGATTTGCCAGCGTGGC**
**CCTCCCTCCAGCAGCACCTGCCCTGGGCTGGGCCACGAGAAGCCCTAGGAGCCCCTGGGGACAGACACACAGCC**
**CCTGCCTCTGTAGGAGACTGTCCTGTTCTGTGAGCGCCCTGTCCTCCGACCCGCATGCCCACTCGGGGGCATGCC**
**TAGTCCATGTGCGTAGGGACAGGCCCTCCCTCACCCATCTACCCCCACGGCACTAACCCCTGGCAGCCCTGCCCA**
**GCCTCGAACCCACATGGGGACACAACCGACTCCGGGGACATGCACTCTCGGGCCCTGTGGAGGGACTGGTCCAG**
**ATGCCCACACACACACTCAGCCCAGACCCGTTCAACAAACCCCGCACTGAGGTTGGCCGGCCACACGGCCACCA**
**CACACACACGTGCACGCCTCACACACGGAGCCTCACCCGGGCGAACCGCACA**

**e.** LHA-EEK-J3-HiCH2CH3(**LS**)-**RHA**

ATCTCTTCCTCAGCACCTGAGTTCCTGGGGGGACCATCAGTCTTCCTGTTCCCCCCAAAACCCAAGGACACTCTCAT
GATCTCCCGGACCCCTGAGGTCACGTGCGTGGTGGTGGACGTGAGCCAGGAAGACCCCGAGGTCCAGTTCAACTG
GTACGTGGATGGCGTGGAGGTGCATAATGCCAAGACAAAGCCGCGGGAGGAGCAGTTCAACAGCACGTACCGTGT
GGTCAGCGTCCTCACCGTCCTGCACCAGGACTGGCTGAACGGCAAGGAGTACAAGTGCAAGGTCTCCAACAAAGGC
CTCCCGTCCTCCATCGAGAAAACCATCTCCAAAGCCAAAGGTGGGACCCACGGGGTGCGAGGGCCACATGGACAGA
GGTCAGCTCGGCCCACCCTCTGCCCTGGGAGTGACCGCTGTGCCAACCTCTGTCCCTACAGGGCAGCCCCGAGAG
CCACAGGTGTACACCCTGCCCCCATCCCAGGAGGAGATGACCAAGAACCAGGTCAGCCTGACCTGCCTGGTCAAAG
GCTTCTACCCCAGCGACATCGCCGTGGAGTGGGAGAGCAATGGGCAGCCGGAGAACAACTACAAGACCACGCCTC

CCGTGCTGGACTCCGACGGCTCCTTCTTCCTCTACAGCAGGCTCACCGTGGACAAGAGCAGGTGGCAGGAGGGGA
ATGTCTTCTCATGCTCCGTGATGCATGAGGCTCTGCACAACCACTACACACAGAAGAGCCTCTCCCTGTCATCGATTA
AACCGGTGAGTTTCATGGTTACTTGCCTGAGAAGATTAAAAAAAGTAATGCTACCTTATGAGGGAGAGTCCCAGGGA
CCAAGATAGCAACTGTCATAGCAACCGTCACACTGCTTTGGTCAAGGAGAAGACCCTTTGGGGAACTGAAAACAGAA
CCTTGAGCACATCTGTTGCTTTCGCTCCCATCCTCCTCCAACAGGGCTGGGTGGAGCACTCCACACCCTTTCACCGG
TCGTACGGCTCAGCCAGAGTAAAAATCACACCCATGACCTGGCCACTGAGGGCTTGATCAATTCACTTTGAATTTGG
CATTAAATACCATTAAGGTATATTAACTGATTTTAAAATAAGATATATTCGTGACCATGTTTTTAACTTTCAAAAATGTAG
CTGCCAGTGTGTGATTTTATTTCAGTTGTACAAAATATCTAAACCTATAGCAATGTGATTAATAAAAACTTAAACATATT
TTCCAGTACCTTAATTCTGTGATAGGAAAATTTTAATCTGAGTATTTTAATTTCATAATCTCTAAAATAGTTTAATGATTT
GTCATTGTGTTGCTGTCGTTTACCCCAGCTGATCTCAAAAGTGATATTTAAGGAGATTATTTTGGTCTGCAACAACTTG
ATAGGGCTCAGCCTCTCCCACCCAACGGGTGGAATCCCCCAGAGGGGGATTTCCAAGAGGCCACCTGGCAGTTGCT
GAGGGTCAGAAGTGAAGCTAGCCACTTCCTCTTAGGCAGGTGGCCAAGATTACAGTTGACCCGTACGTGCAGCTGT
GCCCAGCCTGCCCCATCCCCTGCTCATTTGCATGTTCCCAGAGCACAACCTCCTGCCCTGAAGCCTTATTAATAGGC
TGGTCACACTTTGTGCAGGAGTCAGACTCAGTCAGGACACAGCTGGATCCACTAGTCCAGTGTGGTGGAATTCACCA
TGGAGTTTGGGCTGAGCTGGCTTTTTCTTGTGGCTATTTTAAAAGGTGTCCAGTGTGAGGTGCAGCTGGTGGAGTCT
GGGGGAGGCTTGGTACAGGCCGGGGGGTTCCTGAGACTCTCCTGTGAGCTGAGGGGAAGCATCTTTAACCAGTATG
CCATGGCCTGGTTCCGCCAGGCTCCAGGGAAGGAGAGGGAGTTCGTCGCCGGCATGGGCGCCGTGCCCCACTACG
GCGAGTTCGTGAAGGGCCGGTTCACCATCTCCAGAGACAATGCCAAGAGCACGGTGTATCTGCAAATGAGCAGCCT
GAAGCCCGAGGACACGGCCATCTATTTCTGTGCCAGGAGCAAGAGCACCTACATCAGCTACAACAGCAACGGCTAC
GACTACTGGGGCAGGGGAACCCAGGTCACCGTGAGCAGCGAGCCGAAGAGCTGTGACAAAACACATACATGCCCC
CCCTGCCCCGCGCCCGAGCTGCTTGGCGGTCCAAGCGTGTTCCTGTTCCCCCCCAAGCCCAAGGACACCCTGATGA
TCAGCAGAACCCCCGAGGTGACCTGCGTGGTGGTGGACGTGAGCCACGAGGACCCCGAGGTGAAGTTCAACTGGT
ACGTGGACGGCGTGGAGGTGCACAACGCCAAGACCAAGCCTAGAGAGGAGCAGTACAACAGCACCTACAGAGTGG
TGAGCGTGCTGACCGTGCTGCACCAAGACTGGCTGAACGGCAAGGAGTACAAGTGCAAGGTGTCCAACAAGGCCCT
GCCCGCCCCCATCGAGAAGACCATCAGCAAGGCCAAAGGGCAGCCTAGAGAGCCCCAAGTGTACACCCTGCCCCC
TAGCAGAGACGAGCTGACCAAGAACCAAGTGAGCCTGACCTGCCTGGTGAAGGGCTTCTACCCTAGCGACATCGCC
GTGGAGTGGGAGAGCAACGGGCAGCCCGAGAACAACTACAAGACCACCCCCCCCGTGCTGGACAGCGACGGCAGC
TTCTTCCTGTACAGCAAGCTGACCGTGGACAAGAGCAGATGGCAGCAAGGCAACGTGTTCAGCTGCAGCGTG**CTG**C
ACGAGGCCCTGCAC**TCC**CACTACACACAGAAGAGCCTGAGCCTGAGC**CCGGGTAAATGAGTGCCAGGGCCGGCAA**
**GCCCCCGCTCCCCGGGCTCTCGGGGTCGCGCGAGGATGCTTGGCACGTACCCCGTGTACATACTTCCCGGGCGC**
**CCAGCATGGAAATAAAGCACCCAGCGCTGCCCTGGGCCCCTGCGAGACTGTGATGGTTCTTTCCACGGGTCAGGC**
**CGAGTCTGAGGCCTGAGTGGCATGAGGGAGGCAGAGCGGGTCCCACTGTCCCCACACTGGCCCAGGCTGTGCAG**
**GTGTGCCTGGGCCGCCTAGGGTGGGGCTCAGCCAGGGGCTGCCCTCGGCAGGGTGGGGGATTTGCCAGCGTGGC**
**CCTCCCTCCAGCAGCACCTGCCCTGGGCTGGGCCACGAGAAGCCCTAGGAGCCCCTGGGGACAGACACACAGCC**
**CCTGCCTCTGTAGGAGACTGTCCTGTTCTGTGAGCGCCCTGTCCTCCGACCCGCATGCCCACTCGGGGGCATGCC**
**TAGTCCATGTGCGTAGGGACAGGCCCTCCCTCACCCATCTACCCCCACGGCACTAACCCCTGGCAGCCCTGCCCA**
**GCCTCGAACCCACATGGGGACACAACCGACTCCGGGGACATGCACTCTCGGGCCCTGTGGAGGGACTGGTCCAG**
**ATGCCCACACACACACTCAGCCCAGACCCGTTCAACAAACCCCGCACTGAGGTTGGCCGGCCACACGGCCACCA**
**CACACACACGTGCACGCCTCACACACGGAGCCTCACCCGGGCGAACCGCACA**

**f.** LHA-EEK-J3-HiCH2CH3(**DKD**)-**RHA**

ATCTCTTCCTCAGCACCTGAGTTCCTGGGGGGACCATCAGTCTTCCTGTTCCCCCCAAAACCCAAGGACACTCTCAT
GATCTCCCGGACCCCTGAGGTCACGTGCGTGGTGGTGGACGTGAGCCAGGAAGACCCCGAGGTCCAGTTCAACTG
GTACGTGGATGGCGTGGAGGTGCATAATGCCAAGACAAAGCCGCGGGAGGAGCAGTTCAACAGCACGTACCGTGT
GGTCAGCGTCCTCACCGTCCTGCACCAGGACTGGCTGAACGGCAAGGAGTACAAGTGCAAGGTCTCCAACAAAGGC
CTCCCGTCCTCCATCGAGAAAACCATCTCCAAAGCCAAAGGTGGGACCCACGGGGTGCGAGGGCCACATGGACAGA
GGTCAGCTCGGCCCACCCTCTGCCCTGGGAGTGACCGCTGTGCCAACCTCTGTCCCTACAGGGCAGCCCCGAGAG
CCACAGGTGTACACCCTGCCCCCATCCCAGGAGGAGATGACCAAGAACCAGGTCAGCCTGACCTGCCTGGTCAAAG
GCTTCTACCCCAGCGACATCGCCGTGGAGTGGGAGAGCAATGGGCAGCCGGAGAACAACTACAAGACCACGCCTC
CCGTGCTGGACTCCGACGGCTCCTTCTTCCTCTACAGCAGGCTCACCGTGGACAAGAGCAGGTGGCAGGAGGGGA
ATGTCTTCTCATGCTCCGTGATGCATGAGGCTCTGCACAACCACTACACACAGAAGAGCCTCTCCCTGTCATCGATTA
AACCGGTGAGTTTCATGGTTACTTGCCTGAGAAGATTAAAAAAAGTAATGCTACCTTATGAGGGAGAGTCCCAGGGA
CCAAGATAGCAACTGTCATAGCAACCGTCACACTGCTTTGGTCAAGGAGAAGACCCTTTGGGGAACTGAAAACAGAA

CCTTGAGCACATCTGTTGCTTTCGCTCCCATCCTCCTCCAACAGGGCTGGGTGGAGCACTCCACACCCTTTCACCGG
TCGTACGGCTCAGCCAGAGTAAAAATCACACCCATGACCTGGCCACTGAGGGCTTGATCAATTCACTTTGAATTTGG
CATTAAATACCATTAAGGTATATTAACTGATTTTAAAATAAGATATATTCGTGACCATGTTTTTAACTTTCAAAAATGTAG
CTGCCAGTGTGTGATTTTATTTCAGTTGTACAAAATATCTAAACCTATAGCAATGTGATTAATAAAAACTTAAACATATT
TTCCAGTACCTTAATTCTGTGATAGGAAAATTTTAATCTGAGTATTTTAATTTCATAATCTCTAAAATAGTTTAATGATTT
GTCATTGTGTTGCTGTCGTTTACCCCAGCTGATCTCAAAAGTGATATTTAAGGAGATTATTTTGGTCTGCAACAACTTG
ATAGGGCTCAGCCTCTCCCACCCAACGGGTGGAATCCCCCAGAGGGGGATTTCCAAGAGGCCACCTGGCAGTTGCT
GAGGGTCAGAAGTGAAGCTAGCCACTTCCTCTTAGGCAGGTGGCCAAGATTACAGTTGACCCGTACGTGCAGCTGT
GCCCAGCCTGCCCCATCCCCTGCTCATTTGCATGTTCCCAGAGCACAACCTCCTGCCCTGAAGCCTTATTAATAGGC
TGGTCACACTTTGTGCAGGAGTCAGACTCAGTCAGGACACAGCTGGATCCACTAGTCCAGTGTGGTGGAATTCACCA
TGGAGTTTGGGCTGAGCTGGCTTTTTCTTGTGGCTATTTTAAAAGGTGTCCAGTGTGAGGTGCAGCTGGTGGAGTCT
GGGGGAGGCTTGGTACAGGCCGGGGGGTTCCTGAGACTCTCCTGTGAGCTGAGGGGAAGCATCTTTAACCAGTATG
CCATGGCCTGGTTCCGCCAGGCTCCAGGGAAGGAGAGGGAGTTCGTCGCCGGCATGGGCGCCGTGCCCCACTACG
GCGAGTTCGTGAAGGGCCGGTTCACCATCTCCAGAGACAATGCCAAGAGCACGGTGTATCTGCAAATGAGCAGCCT
GAAGCCCGAGGACACGGCCATCTATTTCTGTGCCAGGAGCAAGAGCACCTACATCAGCTACAACAGCAACGGCTAC
GACTACTGGGGCAGGGGAACCCAGGTCACCGTGAGCAGCGAGCCGAAGAGCTGTGACAAAACACATACATGCCCC
CCCTGCCCCGCGCCCGAGCTGCTTGGCGGTCCAAGCGTGTTCCTGTTCCCCCCCAAGCCCAAGGACACCCTGATGA
TCAGCAGAACCCCCGAGGTGACCTGCGTGGTGGTGGACGTGAGCCACGAGGACCCCGAGGTGAAGTTCAACTGGT
ACGTGGACGGCGTGGAGGTGCACAACGCCAAGACCAAGCCTAGAGAGGAGCAGTACAACAGCACCTACAGAGTGG
TGAGCGTGCTGACCGTGCTGCACCAAGACTGGCTGAACGGCAAGGAGTACAAGTGCAAGGTGTCCAACAAGGCCCT
GCCCGCCCCCATCGAGAAGACCATCAGCAAGGCCAAAGGGCAGCCTAGAGAGCCCCAAGTGTACACCCTGCCCCC
TAGCAGAGACGAGCTGACCAAGAACCAAGTGAGCCTGACCTGCCTGGTGAAGGGCTTCTACCCTAGCGACATCGCC
GTGGAGTGGGAGAGCAACGGGCAGCCCGAGAACAACTAC**GAC**ACCACCCCCCCCGTGCTG**AAG**AGCGACGGCAG
CTTCTTCCTGTACAGC**GAC**CTGACCGTGGACAAGAGCAGATGGCAGCAAGGCAACGTGTTCAGCTGCAGCGTGATG
CACGAGGCCCTGCACAACCACTACACACAGAAGAGCCTGAGCCTGAGC**CCGGGTAAATGAGTGCCAGGGCCGGCA**
**AGCCCCCGCTCCCCGGGCTCTCGGGGTCGCGCGAGGATGCTTGGCACGTACCCCGTGTACATACTTCCCGGGCG**
**CCCAGCATGGAAATAAAGCACCCAGCGCTGCCCTGGGCCCCTGCGAGACTGTGATGGTTCTTTCCACGGGTCAGG**
**CCGAGTCTGAGGCCTGAGTGGCATGAGGGAGGCAGAGCGGGTCCCACTGTCCCCACACTGGCCCAGGCTGTGCA**
**GGTGTGCCTGGGCCGCCTAGGGTGGGGCTCAGCCAGGGGCTGCCCTCGGCAGGGTGGGGGATTTGCCAGCGTGG**
**CCCTCCCTCCAGCAGCACCTGCCCTGGGCTGGGCCACGAGAAGCCCTAGGAGCCCCTGGGGACAGACACACAGC**
**CCCTGCCTCTGTAGGAGACTGTCCTGTTCTGTGAGCGCCCTGTCCTCCGACCCGCATGCCCACTCGGGGGCATGC**
**CTAGTCCATGTGCGTAGGGACAGGCCCTCCCTCACCCATCTACCCCCACGGCACTAACCCCTGGCAGCCCTGCCC**
**AGCCTCGAACCCACATGGGGACACAACCGACTCCGGGGACATGCACTCTCGGGCCCTGTGGAGGGACTGGTCCA**
**GATGCCCACACACACACTCAGCCCAGACCCGTTCAACAAACCCCGCACTGAGGTTGGCCGGCCACACGGCCACC**
**ACACACACACGTGCACGCCTCACACACGGAGCCTCACCCGGGCGAACCGCACA**

**g.** LHA-EEK-J3-HiCH2CH3-modSD-mGFP-**RHA**

ATCTCTTCCTCAGCACCTGAGTTCCTGGGGGGACCATCAGTCTTCCTGTTCCCCCCAAAACCCAAGGACACTCTCAT
GATCTCCCGGACCCCTGAGGTCACGTGCGTGGTGGTGGACGTGAGCCAGGAAGACCCCGAGGTCCAGTTCAACTG
GTACGTGGATGGCGTGGAGGTGCATAATGCCAAGACAAAGCCGCGGGAGGAGCAGTTCAACAGCACGTACCGTGT
GGTCAGCGTCCTCACCGTCCTGCACCAGGACTGGCTGAACGGCAAGGAGTACAAGTGCAAGGTCTCCAACAAAGGC
CTCCCGTCCTCCATCGAGAAAACCATCTCCAAAGCCAAAGGTGGGACCCACGGGGTGCGAGGGCCACATGGACAGA
GGTCAGCTCGGCCCACCCTCTGCCCTGGGAGTGACCGCTGTGCCAACCTCTGTCCCTACAGGGCAGCCCCGAGAG
CCACAGGTGTACACCCTGCCCCCATCCCAGGAGGAGATGACCAAGAACCAGGTCAGCCTGACCTGCCTGGTCAAAG
GCTTCTACCCCAGCGACATCGCCGTGGAGTGGGAGAGCAATGGGCAGCCGGAGAACAACTACAAGACCACGCCTC
CCGTGCTGGACTCCGACGGCTCCTTCTTCCTCTACAGCAGGCTCACCGTGGACAAGAGCAGGTGGCAGGAGGGGA
ATGTCTTCTCATGCTCCGTGATGCATGAGGCTCTGCACAACCACTACACACAGAAGAGCCTCTCCCTGTCATCGATTA
AACCGGTGAGTTTCATGGTTACTTGCCTGAGAAGATTAAAAAAAGTAATGCTACCTTATGAGGGAGAGTCCCAGGGA
CCAAGATAGCAACTGTCATAGCAACCGTCACACTGCTTTGGTCAAGGAGAAGACCCTTTGGGGAACTGAAAACAGAA
CCTTGAGCACATCTGTTGCTTTCGCTCCCATCCTCCTCCAACAGGGCTGGGTGGAGCACTCCACACCCTTTCACCGG
TCGTACGGCTCAGCCAGAGTAAAAATCACACCCATGACCTGGCCACTGAGGGCTTGATCAATTCACTTTGAATTTGG
CATTAAATACCATTAAGGTATATTAACTGATTTTAAAATAAGATATATTCGTGACCATGTTTTTAACTTTCAAAAATGTAG
CTGCCAGTGTGTGATTTTATTTCAGTTGTACAAAATATCTAAACCTATAGCAATGTGATTAATAAAAACTTAAACATATT

TTCCAGTACCTTAATTCTGTGATAGGAAAATTTTAATCTGAGTATTTTAATTTCATAATCTCTAAAATAGTTTAATGATTT
GTCATTGTGTTGCTGTCGTTTACCCCAGCTGATCTCAAAAGTGATATTTAAGGAGATTATTTTGGTCTGCAACAACTTG
ATAGGGCTCAGCCTCTCCCACCCAACGGGTGGAATCCCCCAGAGGGGGATTTCCAAGAGGCCACCTGGCAGTTGCT
GAGGGTCAGAAGTGAAGCTAGCCACTTCCTCTTAGGCAGGTGGCCAAGATTACAGTTGACCCGTACGTGCAGCTGT
GCCCAGCCTGCCCCATCCCCTGCTCATTTGCATGTTCCCAGAGCACAACCTCCTGCCCTGAAGCCTTATTAATAGGC
TGGTCACACTTTGTGCAGGAGTCAGACTCAGTCAGGACACAGCTGGATCCACTAGTCCAGTGTGGTGGAATTCACCA
TGGAGTTTGGGCTGAGCTGGCTTTTTCTTGTGGCTATTTTAAAAGGTGTCCAGTGTGAGGTGCAGCTGGTGGAGTCT
GGGGGAGGCTTGGTACAGGCCGGGGGGTTCCTGAGACTCTCCTGTGAGCTGAGGGGAAGCATCTTTAACCAGTATG
CCATGGCCTGGTTCCGCCAGGCTCCAGGGAAGGAGAGGGAGTTCGTCGCCGGCATGGGCGCCGTGCCCCACTACG
GCGAGTTCGTGAAGGGCCGGTTCACCATCTCCAGAGACAATGCCAAGAGCACGGTGTATCTGCAAATGAGCAGCCT
GAAGCCCGAGGACACGGCCATCTATTTCTGTGCCAGGAGCAAGAGCACCTACATCAGCTACAACAGCAACGGCTAC
GACTACTGGGGCAGGGGAACCCAGGTCACCGTGAGCAGCGAGCCGAAGAGCTGTGACAAAACACATACATGCCCC
CCCTGCCCCGCGCCCGAGCTGCTTGGCGGTCCAAGCGTGTTCCTGTTCCCCCCCAAGCCCAAGGACACCCTGATGA
TCAGCAGAACCCCCGAGGTGACCTGCGTGGTGGTGGACGTGAGCCACGAGGACCCCGAGGTGAAGTTCAACTGGT
ACGTGGACGGCGTGGAGGTGCACAACGCCAAGACCAAGCCTAGAGAGGAGCAGTACAACAGCACCTACAGAGTGG
TGAGCGTGCTGACCGTGCTGCACCAAGACTGGCTGAACGGCAAGGAGTACAAGTGCAAGGTGTCCAACAAGGCCCT
GCCCGCCCCCATCGAGAAGACCATCAGCAAGGCCAAAGGGCAGCCTAGAGAGCCCCAAGTGTACACCCTGCCCCC
TAGCAGAGACGAGCTGACCAAGAACCAAGTGAGCCTGACCTGCCTGGTGAAGGGCTTCTACCCTAGCGACATCGCC
GTGGAGTGGGAGAGCAACGGGCAGCCCGAGAACAACTACAAGACCACCCCCCCCGTGCTGGACAGCGACGGCAGC
TTCTTCCTGTACAGCAAGCTGACCGTGGACAAGAGCAGATGGCAGCAAGGCAACGTGTTCAGCTGCAGCGTGATGC
ACGAGGCCCTGCACAACCACTACACACAGAAGAGCCTGAGCCTGAGCCCaGGTAAA*TCCGGTGGCGGT*GTGAGCAA
GGGCGAGGAGCTGTTCACCGGGGTGGTGCCCATCCTGGTCGAGCTGGACGGCGACGTAAACGGCCACAAGTTCAG
CGTGTCCGGCGAGGGCGAGGGCGATGCCACCTACGGCAAGCTGACCCTGAAGTTCATCTGCACCACCGGCAAGCT
GCCCGTGCCCTGGCCCACCCTCGTGACCACCCTGACCTACGGCGTGCAGTGCTTCAGTCGCTACCCCGACCACATG
AAGCAGCACGACTTCTTCAAGTCCGCCATGCCCGAAGGCTACGTCCAGGAGCGCACCATCTTCTTCAAGGACGACG
GCAACTACAAGACCCGCGCCGAGGTGAAGTTCGAGGGCGACACCCTGGTGAACCGCATCGAGCTGAAGGGCATCG
ACTTCAAGGAGGACGGCAACATCCTGGGGCACAAGCTGGAGTACAACTACAACAGCCACAACGTCTATATCATGGCC
GACAAGCAGAAGAACGGCATCAAGGTGAACTTCAAGATCCGCCACAACATCGAGGACGGCAGCGTGCAGCTCGCCG
ACCACTACCAGCAGAACACCCCCATCGGCGACGGCCCCGTGCTGCTGCCTGACAACCACTACCTGAGCACCCAGTC
CGCCCTGAGCAAAGACCCCAACGAGAAGCGCGATCACATGGTCCTGCTGGAGTTCGTGACCGCCGCCGGGATCACT
CTCGGCATGGACGAGCTGTACAAa**TGAGTGCCAGGGCCGGCAAGCCCCCGCTCCCCGGGCTCTCGGGGTCGCGC**
**GAGGATGCTTGGCACGTACCCCGTGTACATACTTCCCGGGCGCCCAGCATGGAAATAAAGCACCCAGCGCTGCCC**
**TGGGCCCCTGCGAGACTGTGATGGTTCTTTCCACGGGTCAGGCCGAGTCTGAGGCCTGAGTGGCATGAGGGAGG**
**CAGAGCGGGTCCCACTGTCCCCACACTGGCCCAGGCTGTGCAGGTGTGCCTGGGCCGCCTAGGGTGGGGCTCAG**
**CCAGGGGCTGCCCTCGGCAGGGTGGGGGATTTGCCAGCGTGGCCCTCCCTCCAGCAGCACCTGCCCTGGGCTGG**
**GCCACGAGAAGCCCTAGGAGCCCCTGGGGACAGACACACAGCCCCTGCCTCTGTAGGAGACTGTCCTGTTCTGTG**
**AGCGCCCTGTCCTCCGACCCGCATGCCCACTCGGGGGCATGCCTAGTCCATGTGCGTAGGGACAGGCCCTCCCTC**
**ACCCATCTACCCCCACGGCACTAACCCCTGGCAGCCCTGCCCAGCCTCGAACCCACATGGGGACACAACCGACTC**
**CGGGGACATGCACTCTCGGGCCCTGTGGAGGGACTGGTCCAGATGCCCACACACACACTCAGCCCAGACCCGTT**
**CAACAAACCCCGCACTGAGGTTGGCCGGCCACACGGCCACCACACACACACGTGCACGCCTCACACACGGAGCC**
**TCACCCGGGCGAACCGCACA**

**h.** LHA-EEK-CD4d1d2-HiCH2CH3-R5m-**RHA**

ATCTCTTCCTCAGCACCTGAGTTCCTGGGGGGACCATCAGTCTTCCTGTTCCCCCCAAAACCCAAGGACACTCTCAT
GATCTCCCGGACCCCTGAGGTCACGTGCGTGGTGGTGGACGTGAGCCAGGAAGACCCCGAGGTCCAGTTCAACTG
GTACGTGGATGGCGTGGAGGTGCATAATGCCAAGACAAAGCCGCGGGAGGAGCAGTTCAACAGCACGTACCGTGT
GGTCAGCGTCCTCACCGTCCTGCACCAGGACTGGCTGAACGGCAAGGAGTACAAGTGCAAGGTCTCCAACAAAGGC
CTCCCGTCCTCCATCGAGAAAACCATCTCCAAAGCCAAAGGTGGGACCCACGGGGTGCGAGGGCCACATGGACAGA
GGTCAGCTCGGCCCACCCTCTGCCCTGGGAGTGACCGCTGTGCCAACCTCTGTCCCTACAGGGCAGCCCCGAGAG
CCACAGGTGTACACCCTGCCCCCATCCCAGGAGGAGATGACCAAGAACCAGGTCAGCCTGACCTGCCTGGTCAAAG
GCTTCTACCCCAGCGACATCGCCGTGGAGTGGGAGAGCAATGGGCAGCCGGAGAACAACTACAAGACCACGCCTC
CCGTGCTGGACTCCGACGGCTCCTTCTTCCTCTACAGCAGGCTCACCGTGGACAAGAGCAGGTGGCAGGAGGGGA
ATGTCTTCTCATGCTCCGTGATGCATGAGGCTCTGCACAACCACTACACACAGAAGAGCCTCTCCCTGTCATCGATTA

AACCGGTGAGTTTCATGGTTACTTGCCTGAGAAGATTAAAAAAAGTAATGCTACCTTATGAGGGAGAGTCCCAGGGA
CCAAGATAGCAACTGTCATAGCAACCGTCACACTGCTTTGGTCAAGGAGAAGACCCTTTGGGGAACTGAAAACAGAA
CCTTGAGCACATCTGTTGCTTTCGCTCCCATCCTCCTCCAACAGGGCTGGGTGGAGCACTCCACACCCTTTCACCGG
TCGTACGGCTCAGCCAGAGTAAAAATCACACCCATGACCTGGCCACTGAGGGCTTGATCAATTCACTTTGAATTTGG
CATTAAATACCATTAAGGTATATTAACTGATTTTAAAATAAGATATATTCGTGACCATGTTTTTAACTTTCAAAAATGTAG
CTGCCAGTGTGTGATTTTATTTCAGTTGTACAAAATATCTAAACCTATAGCAATGTGATTAATAAAAACTTAAACATATT
TTCCAGTACCTTAATTCTGTGATAGGAAAATTTTAATCTGAGTATTTTAATTTCATAATCTCTAAAATAGTTTAATGATTT
GTCATTGTGTTGCTGTCGTTTACCCCAGCTGATCTCAAAAGTGATATTTAAGGAGATTATTTTGGTCTGCAACAACTTG
ATAGGGCTCAGCCTCTCCCACCCAACGGGTGGAATCCCCCAGAGGGGGATTTCCAAGAGGCCACCTGGCAGTTGCT
GAGGGTCAGAAGTGAAGCTAGCCACTTCCTCTTAGGCAGGTGGCCAAGATTACAGTTGACCCGTACGTGCAGCTGT
GCCCAGCCTGCCCCATCCCCTGCTCATTTGCATGTTCCCAGAGCACAACCTCCTGCCCTGAAGCCTTATTAATAGGC
TGGTCACACTTTGTGCAGGAGTCAGACTCAGTCAGGACACAGCTGGATCCACTAGTCCAGTGTGGTGGAATTCACCA
TGCCCATGGGGTCTCTGCAACCGCTGGCCACCTTGTACCTGCTGGGGATGCTGGTCGCTTCCGTGCTAGCGAAGAA
GGTGGTGCTGGGGAAGAAGGGCGACACCGTGGAGCTGACCTGCACCGCCTCCCAGAAGAAGAGCATCCAGTTCCA
CTGGAAGAACAGCAACCAGATCAAGATCCTGGGAAATCAGGGGAGCTTCCTGACCAAAGGCCCCTCTAAACTGAAC
GACCGGGCTGACTCCCGCCGATCCCTGTGGGATCAGGGCAACTTCCCTTTGATCATCAAAAACCTGAAGATCGAGG
ACTCCGACACCTACATCTGCGAGGTGGAGGATCAAAAGGAGGAGGTGCAACTGCTGGTGTTCGGGCTGACCGCTAA
CAGCGATACCCACCTGCTGCAAGGCCAGTCCCTGACCCTGACCCTGGAGAGCCCACCAGGTTCCAGCCCTTCCGTG
CAGTGCCGGTCTCCCGGGGGCAAGAATATCCAGGGAGGCAAGACCCTGTCTGTGTCCCAGCTGGAGCTCCAGGAC
AGCGGGACCTGGACCTGTACCGTGCTGCAGGACCAGAAGACCGTGGAGTTCAAGATCGACATCGTGGTGCTGGCTG
AGCCGAAGAGCTGTGACAAAACACATACATGCCCCCCCTGCCCCGCGCCCGAGCTGCTTGGCGGTCCAAGCGTGTT
CCTGTTCCCCCCCAAGCCCAAGGACACCCTGATGATCAGCAGAACCCCCGAGGTGACCTGCGTGGTGGTGGACGTG
AGCCACGAGGACCCCGAGGTGAAGTTCAACTGGTACGTGGACGGCGTGGAGGTGCACAACGCCAAGACCAAGCCT
AGAGAGGAGCAGTACAACAGCACCTACAGAGTGGTGAGCGTGCTGACCGTGCTGCACCAAGACTGGCTGAACGGCA
AGGAGTACAAGTGCAAGGTGTCCAACAAGGCCCTGCCCGCCCCCATCGAGAAGACCATCAGCAAGGCCAAAGGGCA
GCCTAGAGAGCCCCAAGTGTACACCCTGCCCCCTAGCAGAGACGAGCTGACCAAGAACCAAGTGAGCCTGACCTGC
CTGGTGAAGGGCTTCTACCCTAGCGACATCGCCGTGGAGTGGGAGAGCAACGGGCAGCCCGAGAACAACTACAAG
ACCACCCCCCCCGTGCTGGACAGCGACGGCAGCTTCTTCCTGTACAGCAAGCTGACCGTGGACAAGAGCAGATGGC
AGCAAGGCAACGTGTTCAGCTGCAGCGTGATGCACGAGGCCCTGCACAACCACTACACACAGAAGAGCCTGAGCCT
GAGCCCGGGTAAA*TCCGGTGGCGGT*GGCGATTATTACGATTACGACGGTGGCTATTACTATGATGGCGAC**TGAGTG**
**CCAGGGCCGGCAAGCCCCCGCTCCCCGGGCTCTCGGGGTCGCGCGAGGATGCTTGGCACGTACCCCGTGTACAT**
**ACTTCCCGGGCGCCCAGCATGGAAATAAAGCACCCAGCGCTGCCCTGGGCCCCTGCGAGACTGTGATGGTTCTTT**
**CCACGGGTCAGGCCGAGTCTGAGGCCTGAGTGGCATGAGGGAGGCAGAGCGGGTCCCACTGTCCCCACACTGGC**
**CCAGGCTGTGCAGGTGTGCCTGGGCCGCCTAGGGTGGGGCTCAGCCAGGGGCTGCCCTCGGCAGGGTGGGGGA**
**TTTGCCAGCGTGGCCCTCCCTCCAGCAGCACCTGCCCTGGGCTGGGCCACGAGAAGCCCTAGGAGCCCCTGGGG**
**ACAGACACACAGCCCCTGCCTCTGTAGGAGACTGTCCTGTTCTGTGAGCGCCCTGTCCTCCGACCCGCATGCCCA**
**CTCGGGGGCATGCCTAGTCCATGTGCGTAGGGACAGGCCCTCCCTCACCCATCTACCCCCACGGCACTAACCCCT**
**GGCAGCCCTGCCCAGCCTCGAACCCACATGGGGACACAACCGACTCCGGGGACATGCACTCTCGGGCCCTGTGG**
**AGGGACTGGTCCAGATGCCCACACACACACTCAGCCCAGACCCGTTCAACAAACCCCGCACTGAGGTTGGCCGG**
**CCACACGGCCACCACACACACACGTGCACGCCTCACACACGGAGCCTCACCCGGGCGAACCGCACA**

**Supplementary Table 3. PCR primer sets.** Primers for detecting on- and off-target editing at (**a**) the *IGHG1* CH2 intron, and (**b**) the *IGHG4* CH3 exon. (**c**) Primer sets for nested in-out PCR, to confirm J3-CH2 insertion at *IGHG1* CH2 intron.

**a.** *For IGHG1 CH2 intron targeting gRNAs*

| Targets | Locus | Primer name | Sequence |
|---|---|---|---|
| On-target | IGHG1 | IGHG1-CH2-Fwd | ACACCTTCTCTCCTCCCAGATTCC |
| | | IGHG1-CH2-Rev | CGGCGATGTCGCTGGGA |
| Off-target | IGHG2 | IGHG2-CH2-Fwd | CGACCCCAAAGGCCAAACTG |
| | | IGHG2-CH2-Rev | GGAGTCCAGCATGGGAGGT |
| | IGHG3 | IGHG3-CH2-Fwd | GTCGGGTGCTGACACATCTG |
| | | IGHG3-CH2-Rev | GAGGCTCTTCTGCGTGAAGC |
| | IGHG4 | IGHG4-CH2-Fwd | GGGGGACCATCAGTCTTCCTG |
| | | IGHG4-CH2-Rev | GGAGCATGAGAAGACATTCCCCTC |
| | IGHGP | IGHGP-CH2-Fwd | ACACATGCCCACCATGTGCAA |
| | | IGHGP-CH2-Rev | GCTATAGAGGAAGAAGGAGCCGTT |
| Sequencing primer | | CH2-Uniseq | TGGAGGTGCATAATGCCAAGAC |

**b.** *For IGHG4 CH3 exon targeting gRNAs*

| Targets | Locus | Primer name | Sequence |
|---|---|---|---|
| Off-target | IGHG1 | IGHG1-CH2-Fwd | ACACCTTCTCTCCTCCCAGATTCC |
| | | IGHG1-end-Rev | CAGGGACGTGACGTGGT |
| | IGHG2 | IGHG2-CH2-Fwd | CGACCCCAAAGGCCAAACTG |
| | | IGHG2-end-Rev | GCCAGTGTGGGGACAGTGGA |
| | IGHG3 | IGHG3-CH2-Fwd | GTCGGGTGCTGACACATCTG |
| | | IGHG3-end-Rev | AGTTGCAGCTCTGGACAGGAAGA |
| On-target | IGHG4 | IGHG4-CH2-Fwd | GGGGGACCATCAGTCTTCCTG |
| | | IGHG4-end-Rev | CAGGAGGATGGTGAAACCCACC |
| Off-target | IGHGP | IGHGP-CH2-Fwd | ACACATGCCCACCATGTGCAA |
| | | IGHGP-end-Rev | AGCATCCTCGTGCGACCG |
| Sequencing primer | | end-Uniseq | CCTGCCTGGTCAAAGGCTTC |

**c.** *For nested in-out PCR*

| Purpose | Primer name | Sequence |
|---|---|---|
| 1st PCR | IGHG1-HA-Fwd2 | GTCAGGGGCTTCAGGGGG |
| | EEK-Beg-Rev | CTTGACCAAAGCAGTGTGACG |
| 2nd PCR | IGHG1-F1 | CCTGGCACCCTCCTCCAA |
| | IO-EEK-In-6 | GACAGTTGCTATCTTGGTCCCTG |
| Sequencing primer | IO-EEK-In-6 | GACAGTTGCTATCTTGGTCCCTG |

**Supplementary Table 4. Flow cytometry staining panels.** (**a**) Panel for tonsil organoid or PBMC edited human B cells expressing J3 HCAb. (**b**) Panel for ADCC assay (**c**) Panel for J3 HCAb or eCD4-Ig edited Raji cells and eCD4-Ig edited PBMC B cells.

(**a**)

| Antibodies | Fluorescence | Source | Cat. num. | Intracellular stain |
|---|---|---|---|---|
| IgG | BV421 | Biolegend | 410704 | Yes |
| His | PE | MIltenyi | 130-120-718 | |
| VHH | APC | GeneScript | A01994 | Yes |
| CD27 | BV510 | BioLegend | 302836 | |
| CD38 | PerCP-eF710 | Thermo Fisher | 46-0389-42 | |
| CD138 | BV711 | Biolegend | 356522 | |
| CD3 | PE-eF610 | Thermo Fisher | 61-0038-42 | |
| CD19 | SB600 | Thermo Fisher | 63-0198-42 | |
| IgM | FITC | BD | 555782 | |
| IgD | PE-Cy7 | BD | 561314 | |
| Viability dye | Near-IR | Thermo Fisher | L34976 | |

(**b**)

| Antibodies | Fluorescence | Source | Cat. num. | Intracellular stain |
|---|---|---|---|---|
| CD107A | Alexa Fluor 647 | BD | 562622 | Yes |
| CD56 | BB-700 | BD | 566400 | |
| p24 | RD1 (PE) | Beckman Coulter | 6604667 | Yes |
| Viability dye | Near-IR | Thermo Fisher | L34976 | |

(**c**)

| Antibodies | Fluorescence | Source | Cat. Num. |
|---|---|---|---|
| His | APC | Miltenyi | 130-119-782 |
| IgG | PE | Biolegend | 410708 |
| Viability dye | Near-IR | Thermo Fisher | L34976 |

**Supplementary Table 5. Sequences of HCAb expression plasmids.** (**a**, **b**) Expression plasmid for HCAb J3-IgG1, with PGK promoter and (**a**) without and (**b**) with LS mutations (bolded and underlined) in IgG1 CH3. (**c**) Expression plasmid for HCAb eCD4-IgG1, with hybrid CMV/R promoter from plasmid CMVR-eCD4-IgG2-v26. A GGGG linker is present between CH3 and the R5m domain (italicized, underlined). (**d**) Expression plasmid for HCAb CD4-IgG1. The sequences are colored to indicate specific regions, as noted in construct name, with multiple cloning sites between segments in black text.

**a.** PGK-J3-HiCH2CH3-polyA

GGGGTTGGGGTTGCGCCTTTTCCAAGGCAGCCCTGGGTTTGCGCAGGGACGCGGCTGCTCTGGGCGTGGTTCCGG
GAAACGCAGCGGCGCCGACCCTGGGTCTCGCACATTCTTCACGTCCGTTCGCAGCGTCACCCGGATCTTCGCCGCT
ACCCTTGTGGGCCCCCCGGCGACGCTTCCTGCTCCGCCCCTAAGTCGGGAAGGTTCCTTGCGGTTCGCGGCGTGC
CGGACGTGACAAACGGAAGCCGCACGTCTCACTAGTACCCTCGCAGACGGACAGCGCCAGGGAGCAATGGCAGCG
CGCCGACCGCGATGGGCTGTGGCCAATAGCGGCTGCTCAGCAGGGCGCGCCGAGAGCAGCGGCCGGGAAGGGGC
GGTGCGGGAGGCGGGGTGTGGGGCGGTAGTGTGGGCCCTGTTCCTGCCCGCGCGGTGTTCCGCATTCTGCAAGCC
TCCGGAGCGCACGTCGGCAGTCGGCTCCCTCGTTGACCGAATCACCGACCTCTCTCCCCAGGGGGATCCACTAGTC
CAGTGTGGTGGAATTCACCATGGAGTTTGGGCTGAGCTGGCTTTTTCTTGTGGCTATTTTAAAAGGTGTCCAGTGTGA
GGTGCAGCTGGTGGAGTCTGGGGGAGGCTTGGTACAGGCCGGGGGGTTCCTGAGACTCTCCTGTGAGCTGAGGGG
AAGCATCTTTAACCAGTATGCCATGGCCTGGTTCCGCCAGGCTCCAGGGAAGGAGAGGGAGTTCGTCGCCGGCATG
GGCGCCGTGCCCCACTACGGCGAGTTCGTGAAGGGCCGGTTCACCATCTCCAGAGACAATGCCAAGAGCACGGTG
TATCTGCAAATGAGCAGCCTGAAGCCCGAGGACACGGCCATCTATTTCTGTGCCAGGAGCAAGAGCACCTACATCAG
CTACAACAGCAACGGCTACGACTACTGGGGCAGGGGAACCCAGGTCACCGTGAGCAGCGAGCCGAAGAGCTGTGA
CAAAACACATACATGCCCCCCCTGCCCCGCGCCCGAGCTGCTTGGCGGTCCAAGCGTGTTCCTGTTCCCCCCCAAG
CCCAAGGACACCCTGATGATCAGCAGAACCCCCGAGGTGACCTGCGTGGTGGTGGACGTGAGCCACGAGGACCCC
GAGGTGAAGTTCAACTGGTACGTGGACGGCGTGGAGGTGCACAACGCCAAGACCAAGCCTAGAGAGGAGCAGTAC
AACAGCACCTACAGAGTGGTGAGCGTGCTGACCGTGCTGCACCAAGACTGGCTGAACGGCAAGGAGTACAAGTGCA
AGGTGTCCAACAAGGCCCTGCCCGCCCCCATCGAGAAGACCATCAGCAAGGCCAAAGGGCAGCCTAGAGAGCCCC
AAGTGTACACCCTGCCCCCTAGCAGAGACGAGCTGACCAAGAACCAAGTGAGCCTGACCTGCCTGGTGAAGGGCTT
CTACCCTAGCGACATCGCCGTGGAGTGGGAGAGCAACGGGCAGCCCGAGAACAACTACAAGACCACCCCCCCCGT
GCTGGACAGCGACGGCAGCTTCTTCCTGTACAGCAAGCTGACCGTGGACAAGAGCAGATGGCAGCAAGGCAACGTG
TTCAGCTGCAGCGTGATGCACGAGGCCCTGCACAACCACTACACACAGAAGAGCCTGAGCCTGAGCCCGGGTAAAT
GAGTCTAGAGGGCCCGTTTAAACCCGCTGATCAGCCTCGA<u>CTGTGCCTTCTAGTTGCCAGCCATCTGTTGTTTGCCC</u>
<u>CTCCCCCGTGCCTTCCTTGACCCTGGAAGGTGCCACTCCCACTGTCCTTTCCTAATAAAATGAGGAAATTGCATCGCA</u>
<u>TTGTCTGAGTAGGTGTCATTCTATTCTGGGGGGTGGGGTGGGGCAGGACAGCAAGGGGGAGGATTGGGAAGACAAT</u>
<u>AGCAGGCATGCTGGGGATGCGGTGGGCTCTATGG</u>

**b.** PGK-J3-HiCH2CH3(**<u>LS</u>**)-polyA

GGGGTTGGGGTTGCGCCTTTTCCAAGGCAGCCCTGGGTTTGCGCAGGGACGCGGCTGCTCTGGGCGTGGTTCCGG
GAAACGCAGCGGCGCCGACCCTGGGTCTCGCACATTCTTCACGTCCGTTCGCAGCGTCACCCGGATCTTCGCCGCT
ACCCTTGTGGGCCCCCCGGCGACGCTTCCTGCTCCGCCCCTAAGTCGGGAAGGTTCCTTGCGGTTCGCGGCGTGC
CGGACGTGACAAACGGAAGCCGCACGTCTCACTAGTACCCTCGCAGACGGACAGCGCCAGGGAGCAATGGCAGCG
CGCCGACCGCGATGGGCTGTGGCCAATAGCGGCTGCTCAGCAGGGCGCGCCGAGAGCAGCGGCCGGGAAGGGGC
GGTGCGGGAGGCGGGGTGTGGGGCGGTAGTGTGGGCCCTGTTCCTGCCCGCGCGGTGTTCCGCATTCTGCAAGCC
TCCGGAGCGCACGTCGGCAGTCGGCTCCCTCGTTGACCGAATCACCGACCTCTCTCCCCAGGGGGATCCACTAGTC
CAGTGTGGTGGAATTCACCATGGAGTTTGGGCTGAGCTGGCTTTTTCTTGTGGCTATTTTAAAAGGTGTCCAGTGTGA
GGTGCAGCTGGTGGAGTCTGGGGGAGGCTTGGTACAGGCCGGGGGGTTCCTGAGACTCTCCTGTGAGCTGAGGGG
AAGCATCTTTAACCAGTATGCCATGGCCTGGTTCCGCCAGGCTCCAGGGAAGGAGAGGGAGTTCGTCGCCGGCATG
GGCGCCGTGCCCCACTACGGCGAGTTCGTGAAGGGCCGGTTCACCATCTCCAGAGACAATGCCAAGAGCACGGTG
TATCTGCAAATGAGCAGCCTGAAGCCCGAGGACACGGCCATCTATTTCTGTGCCAGGAGCAAGAGCACCTACATCAG
CTACAACAGCAACGGCTACGACTACTGGGGCAGGGGAACCCAGGTCACCGTGAGCAGCGAGCCGAAGAGCTGTGA
CAAAACACATACATGCCCCCCCTGCCCCGCGCCCGAGCTGCTTGGCGGTCCAAGCGTGTTCCTGTTCCCCCCCAAG
CCCAAGGACACCCTGATGATCAGCAGAACCCCCGAGGTGACCTGCGTGGTGGTGGACGTGAGCCACGAGGACCCC

GAGGTGAAGTTCAACTGGTACGTGGACGGCGTGGAGGTGCACAACGCCAAGACCAAGCCTAGAGAGGAGCAGTAC
AACAGCACCTACAGAGTGGTGAGCGTGCTGACCGTGCTGCACCAAGACTGGCTGAACGGCAAGGAGTACAAGTGCA
AGGTGTCCAACAAGGCCCTGCCCGCCCCCATCGAGAAGACCATCAGCAAGGCCAAAGGGCAGCCTAGAGAGCCCC
AAGTGTACACCCTGCCCCCTAGCAGAGACGAGCTGACCAAGAACCAAGTGAGCCTGACCTGCCTGGTGAAGGGCTT
CTACCCTAGCGACATCGCCGTGGAGTGGGAGAGCAACGGGCAGCCCGAGAACAACTACAAGACCACCCCCCCCGT
GCTGGACAGCGACGGCAGCTTCTTCCTGTACAGCAAGCTGACCGTGGACAAGAGCAGATGGCAGCAAGGCAACGTG
TTCAGCTGCAGCGTG**CTG**CACGAGGCCCTGCAC**TCC**CACTACACACAGAAGAGCCTGAGCCTGAGCCCGGGTAAAT
GAGTCTAGAGGGCCCGTTTAAACCCGCTGATCAGCCTCGACTGTGCCTTCTAGTTGCCAGCCATCTGTTGTTTGCCC
CTCCCCCGTGCCTTCCTTGACCCTGGAAGGTGCCACTCCCACTGTCCTTTCCTAATAAAATGAGGAAATTGCATCGCA
TTGTCTGAGTAGGTGTCATTCTATTCTGGGGGGTGGGGTGGGGCAGGACAGCAAGGGGGAGGATTGGGAAGACAAT
AGCAGGCATGCTGGGGATGCGGTGGGCTCTATGG

**c.** CMV/R-CD4d1d2-HiCH2CH3-R5m-polyA

GACATTGATTATTGACTAGTTATTAATAGTAATCAATTACGGGGTCATTAGTTCATAGCCCATATATGGAGTTCCGCGT
TACATAACTTACGGTAAATGGCCCGCCTGGCTGACCGCCCAACGACCCCCGCCCATTGACGTCAATAATGACGTATG
TTCCCATAGTAACGCCAATAGGGACTTTCCATTGACGTCAATGGGTGGAGTATTTACGGTAAACTGCCCACTTGGCAG
TACATCAAGTGTATCATATGCCAAGTACGCCCCCTATTGACGTCAATGACGGTAAATGGCCCGCCTGGCATTATGCC
CAGTACATGACCTTATGGGACTTTCCTACTTGGCAGTACATCTACGTATTAGTCATCGCTATTACCATGGTGATGCGG
TTTTGGCAGTACATCAATGGGCGTGGATAGCGGTTTGACTCACGGGGATTTCCAAGTCTCCACCCCATTGACGTCAA
TGGGAGTTTGTTTTGGCACCAAAATCAACGGGACTTTCCAAAATGTCGTAACAACTCCGCCCCATTGACGCAAATGG
GCGGTAGGCGTGTACGGTGGGAGGTCTATATAAGCAGAGCTCGTTTAGTGAACCGTCAGATCGCCTGGAGACGCCA
TCCACGCTGTTTTGACCTCCATAGAAGACACCGGGACCGATCCAGCCTCCATCGGCTCGCATCTCTCCTTCACGCGC
CCGCCGCCCTACCTGAGGCCGCCATCCACGCCGGTTGAGTCGCGTTCTGCCGCCTCCCGCCTGTGGTGCCTCCTG
AACTGCGTCCGCCGTCTAGGTAAGTTTAAAGCTCAGGTCGAGACCGGGCCTTTGTCCGGCGCTCCCTTGGAGCCTA
CCTAGACTCAGCCGGCTCTCCACGCTTTGCCTGACCCTGCTTGCTCAACTCTAGTTAACGGTGGAGGGCAGTGTAGT
CTGAGCAGTACTCGTTGCTGCCGCGCGCGCCACCAGACATAATAGCTGACAGACTAACAGACTGTTCCTTTCCATGG
GTCTTTTCTGCAGTCACCGTCGTCGACACGTGTGATCAGATATCGCGGCCGCTCTAGACCACCATGCCCATGGGGTC
TCTGCAACCGCTGGCCACCTTGTACCTGCTGGGGATGCTGGTCGCTTCCGTGCTAGCGAAGAAGGTGGTGCTGGGG
AAGAAGGGCGACACCGTGGAGCTGACCTGCACCGCCTCCCAGAAGAAGAGCATCCAGTTCCACTGGAAGAACAGCA
ACCAGATCAAGATCCTGGGAAATCAGGGGAGCTTCCTGACCAAAGGCCCCTCTAAACTGAACGACCGGGCTGACTC
CCGCCGATCCCTGTGGGATCAGGGCAACTTCCCTTTGATCATCAAAAACCTGAAGATCGAGGACTCCGACACCTACA
TCTGCGAGGTGGAGGATCAAAAGGAGGAGGTGCAACTGCTGGTGTTCGGGCTGACCGCTAACAGCGATACCCACCT
GCTGCAAGGCCAGTCCCTGACCCTGACCCTGGAGAGCCCACCAGGTTCCAGCCCTTCCGTGCAGTGCCGGTCTCCC
GGGGGCAAGAATATCCAGGGAGGCAAGACCCTGTCTGTGTCCCAGCTGGAGCTCCAGGACAGCGGGACCTGGACC
TGTACCGTGCTGCAGGACCAGAAGACCGTGGAGTTCAAGATCGACATCGTGGTGCTGGCTGAGCCCAAATCTTGTG
ACAAAACTCACACATGCCCACCGTGCCCAGCACCTGAACTCCTGGGGGGACCGTCAGTCTTCCTCTTCCCCCCAAAA
CCCAAGGACACCCTCATGATCTCCCGGACCCCTGAGGTCACATGCGTGGTGGTGGACGTGAGCCACGAAGACCCTG
AGGTCAAGTTCAACTGGTACGTGGACGGCGTGGAGGTGCATAATGCCAAGACAAAGCCGCGGGAGGAGCAGTACAA
CAGCACGTACCGTGTGGTCAGCGTCCTCACCGTCCTGCACCAGGACTGGCTGAATGGCAAGGAGTACAAGTGCAAG
GTCTCCAACAAAGCCCTCCCAGCCCCCATCGAGAAAACCATCTCCAAAGCCAAAGGGCAGCCCCGAGAACCACAGG
TGTACACCCTGCCCCCATCCCGGGATGAGCTGACCAAGAACCAGGTCAGCCTGACCTGCCTGGTCAAAGGCTTCTA
TCCCAGCGACATCGCCGTGGAGTGGGAGAGCAATGGGCAGCCGGAGAACAACTACAAGACCACGCCTCCCGTGCT
GGACTCCGATGGCAGCTTCTTCCTCTATTCAAAGCTGACAGTGGACAAATCCAGATGGCAGCAGGGGAACGTCTTTA
GCTGCTCTGTGATGCACGAGGCCCTGCACAATCATTACACCCAGAAGAGTCTCTCACTGTCCCCCGGAAAG*GGCGG*
*TGGCGGT*GGCGATTATTACGATTACGACGGTGGCTATTACTATGATGGCGACTGAGGATCCAGATCTGCTGTGCCTT
CTAGTTGCCAGCCATCTGTTGTTTGCCCCTCCCCCGTGCCTTCCTTGACCCTGGAAGGTGCCACTCCCACTGTCCTT
TCCTAATAAAATGAGGAAATTGCATCGCATTGTCTGAGTAGGTGTCATTCTATTCTGGGGGGTGGGGTGGGGCAGGA
CAGCAAGGGGGAGGATTGGGAAGACAATAGCAGGCATGCTGGGGATGCGGTGGGCTCTATGGG

**d.** CMV/R-CD4dd2-HiCH2CH3-polyA

GACATTGATTATTGACTAGTTATTAATAGTAATCAATTACGGGGTCATTAGTTCATAGCCCATATATGGAGTTCCGCGT
TACATAACTTACGGTAAATGGCCCGCCTGGCTGACCGCCCAACGACCCCCGCCCATTGACGTCAATAATGACGTATG

TTCCCATAGTAACGCCAATAGGGACTTTCCATTGACGTCAATGGGTGGAGTATTTACGGTAAACTGCCCACTTGGCAG
TACATCAAGTGTATCATATGCCAAGTACGCCCCCTATTGACGTCAATGACGGTAAATGGCCCGCCTGGCATTATGCC
CAGTACATGACCTTATGGGACTTTCCTACTTGGCAGTACATCTACGTATTAGTCATCGCTATTACCATGGTGATGCGG
TTTTGGCAGTACATCAATGGGCGTGGATAGCGGTTTGACTCACGGGGATTTCCAAGTCTCCACCCCATTGACGTCAA
TGGGAGTTTGTTTTGGCACCAAAATCAACGGGACTTTCCAAAATGTCGTAACAACTCCGCCCCATTGACGCAAATGG
GCGGTAGGCGTGTACGGTGGGAGGTCTATATAAGCAGAGCTCGTTTAGTGAACCGTCAGATCGCCTGGAGACGCCA
TCCACGCTGTTTTGACCTCCATAGAAGACACCGGGACCGATCCAGCCTCCATCGGCTCGCATCTCTCCTTCACGCGC
CCGCCGCCCTACCTGAGGCCGCCATCCACGCCGGTTGAGTCGCGTTCTGCCGCCTCCCGCCTGTGGTGCCTCCTG
AACTGCGTCCGCCGTCTAGGTAAGTTTAAAGCTCAGGTCGAGACCGGGCCTTTGTCCGGCGCTCCCTTGGAGCCTA
CCTAGACTCAGCCGGCTCTCCACGCTTTGCCTGACCCTGCTTGCTCAACTCTAGTTAACGGTGGAGGGCAGTGTAGT
CTGAGCAGTACTCGTTGCTGCCGCGCGCGCCACCAGACATAATAGCTGACAGACTAACAGACTGTTCCTTTCCATGG
GTCTTTTCTGCAGTCACCGTCGTCGACACGTGTGATCAGATATCGCGGCCGCTCTAGACCACCATGCCCATGGGGTC
TCTGCAACCGCTGGCCACCTTGTACCTGCTGGGGATGCTGGTCGCTTCCGTGCTAGCGAAGAAGGTGGTGCTGGGG
AAGAAGGGCGACACCGTGGAGCTGACCTGCACCGCCTCCCAGAAGAAGAGCATCCAGTTCCACTGGAAGAACAGCA
ACCAGATCAAGATCCTGGGAAATCAGGGGAGCTTCCTGACCAAAGGCCCCTCTAAACTGAACGACCGGGCTGACTC
CCGCCGATCCCTGTGGGATCAGGGCAACTTCCCTTTGATCATCAAAAACCTGAAGATCGAGGACTCCGACACCTACA
TCTGCGAGGTGGAGGATCAAAAGGAGGAGGTGCAACTGCTGGTGTTCGGGCTGACCGCTAACAGCGATACCCACCT
GCTGCAAGGCCAGTCCCTGACCCTGACCCTGGAGAGCCCACCAGGTTCCAGCCCTTCCGTGCAGTGCCGGTCTCCC
GGGGGCAAGAATATCCAGGGAGGCAAGACCCTGTCTGTGTCCCAGCTGGAGCTCCAGGACAGCGGGACCTGGACC
TGTACCGTGCTGCAGGACCAGAAGACCGTGGAGTTCAAGATCGACATCGTGGTGCTGGCTGAGCCCAAATCTTGTG
ACAAAACTCACACATGCCCACCGTGCCCAGCACCTGAACTCCTGGGGGGACCGTCAGTCTTCCTCTTCCCCCCAAAA
CCCAAGGACACCCTCATGATCTCCCGGACCCCTGAGGTCACATGCGTGGTGGTGGACGTGAGCCACGAAGACCCTG
AGGTCAAGTTCAACTGGTACGTGGACGGCGTGGAGGTGCATAATGCCAAGACAAAGCCGCGGGAGGAGCAGTACAA
CAGCACGTACCGTGTGGTCAGCGTCCTCACCGTCCTGCACCAGGACTGGCTGAATGGCAAGGAGTACAAGTGCAAG
GTCTCCAACAAAGCCCTCCCAGCCCCCATCGAGAAAACCATCTCCAAAGCCAAAGGGCAGCCCCGAGAACCACAGG
TGTACACCCTGCCCCCATCCCGGGATGAGCTGACCAAGAACCAGGTCAGCCTGACCTGCCTGGTCAAAGGCTTCTA
TCCCAGCGACATCGCCGTGGAGTGGGAGAGCAATGGGCAGCCGGAGAACAACTACAAGACCACGCCTCCCGTGCT
GGACTCCGATGGCAGCTTCTTCCTCTATTCAAAGCTGACAGTGGACAAATCCAGATGGCAGCAGGGGAACGTCTTTA
GCTGCTCTGTGATGCACGAGGCCCTGCACAATCATTACACCCAGAAGAGTCTCTCACTGTCCCCCGGAAAGTGAGGA
TCCAGATCTGCTGTGCCTTCTAGTTGCCAGCCATCTGTTGTTTGCCCCTCCCCCGTGCCTTCCTTGACCCTGGAAGG
TGCCACTCCCACTGTCCTTTCCTAATAAAATGAGGAAATTGCATCGCATTGTCTGAGTAGGTGTCATTCTATTCTGGG
GGGTGGGGTGGGGCAGGACAGCAAGGGGGAGGATTGGGAAGACAATAGCAGGCATGCTGGGGATGCGGTGGGCT
CTATGG

**Supplementary Table 6. Primers and probes used for ddPCR.**

| Detection | Primer/ probe | Sequence |
|---|---|---|
| Total J3 HCAb or eCD4-IgG1 mRNA | dd-optHiCH2-F1 | GGACACCCTGATGATCAGCAGA |
| | dd-optHiCH2-R1 | CCTTTGGCCTTGCTGATGGTC |
| | dd-optHiCH2-probe (HEX) | /5HEX/CAGTACAAC/ZEN/AGCACCTACAGAGTGG/3IABkFQ/ |
| J3 HCAb BCR or eCD4-IgG1 BCR mRNA | dd-optCH3M1-F1 | CTGAGCCTGAGCCCGGAG |
| | dd-G4BCR-R2 | GGCCCCCTGCCTTATCATGT |
| | dd-BCR-probe | /56-FAM/TCTTCTCCT/ZEN/CAGTGGTGGACCTGA/3IABkFQ/ |
| J3 HCAb secreted mRNA | dd-optCH3-F1 | CCTGCCCCCTAGCAGAGAC |
| | dd-optCH3-sec-R1 | GGCACTCATTTACCCGGGCT |
| | dd-sec-probe | /56-FAM/TGACCAAGA/ZEN/ACCAAGTGAGCCTGA/3IABkFQ/ |
| eCD4-IgG1 secreted mRNA | dd-optCH3-F1 | CCTGCCCCCTAGCAGAGAC |
| | dd-CCR5m-R1 | CGTCGTAATCGTAATAATCGCCACC |
| | dd-sec-probe | /56-FAM/TGACCAAGA/ZEN/ACCAAGTGAGCCTGA/3IABkFQ/ |
